# Plastid phylogenomics clarifies broad-level relationships in *Bulbophyllum* (Orchidaceae) and provides insights into range evolution of Australasian section *Adelopetalum*

**DOI:** 10.1101/2022.07.24.500920

**Authors:** Lalita Simpson, Mark A. Clements, Harvey K. Orel, Darren M. Crayn, Katharina Nargar

## Abstract

The hyper diverse orchid genus *Bulbophyllum* is the second largest genus of flowering plants and exhibits a pantropical distribution with a center of diversity in tropical Asia. The only *Bulbophyllum* section with a center of diversity in Australasia is sect. *Adelopetalum*. However, phylogenetic placement, interspecific relationships, and spatio-temporal evolution of the section have remained largely unclear. To infer broad-level relationships within *Bulbophyllum* and interspecific relationships within sect. *Adelopetalum*, a genome skimming dataset was generated for 89 samples, yielding 70 plastid coding regions and the nuclear ribosomal DNA cistron. For 18 additional samples, Sanger data from two plastid loci (*mat*K, *ycf*1) and nuclear ITS were added using a supermatrix approach. The study provided new insights into broad-level relationships in *Bulbophyllum*, including phylogenetic evidence for the non-monophyly of sections *Beccariana, Brachyantha, Brachypus, Cirrhopetaloides, Cirrhopetalum, Desmosanthes, Minutissima, Oxysepala, Polymeres* and *Sestochilos*. Section *Adelopetalum* and sect. *Minutissima s*.*s*. formed a highly supported clade that was resolved in sister group position to the remainder of the genus. Divergence time estimations based on a relaxed molecular clock model placed the origin of *Bulbophyllum* in the early Oligocene (ca. 33.2 Ma) and of sect. *Adelopetalum* in the late Oligocene (ca. 23.6 Ma). Ancestral range estimations based on a BAYAREALIKE model identified the Australian continent as ancestral area of sect. *Adelopetalum*. The section underwent crown diversification during the mid-Miocene to the late Pleistocene, predominantly in continental Australia. At least two independent long-distance dispersal events were inferred eastwards from the Australian continent to New Zealand, and New Caledonia from the early Pliocene onwards, likely mediated by the predominantly westerly winds of the southern hemisphere. Retraction and fragmentation of eastern Australian rainforests from the early Miocene onwards are discussed as likely drivers of lineage divergence within sect. *Adelopetalum*, facilitating allopatric speciation.

## 1 Introduction

The hyper diverse orchid genus *Bulbophyllum* Thouars (Epidendroideae) is the second largest genus of flowering plants with more than 2,100 species and exhibits exceptional morphological and ecological diversity (Frodin, 2004; Pridgeon et al., 2014, WCSP 2022). Species of this predominantly epiphytic genus occur in a wide range of tropical and subtropical habitats, from montane rainforests to dry deciduous forests, savannah woodlands, and rocky fields with shrubby vegetation (Pridgeon et al., 2014). *Bulbophyllum* is distributed pantropically, occupying all botanical continents defined by Brummit (2001) except for Antarctic and Eurasia. The genus is most diverse on the botanical continent of tropical Asia (1562 species), also occurring on the botanical continents of Africa (305), temperate Asia (152), Southern America (88), the Pacific (49), Australasia (Australia and New Zealand; 35), and Northern America (7) (WCSP, 2022). Centres of diversity are found in tropical Asia in the floristic regions of Malesia (667) and Papuasia (656) and in the Afrotropics in the western Indian Ocean region, on the islands of Madagascar, and the Mascarenes (218) (WCSP, 2022).

The high number of species and complex patterns of morphological variation has presented significant challenges for resolving relationships in *Bulbophyllum* and this is reflected in substantial taxonomic revisions that have been proposed. Traditionally, the subtribe Bulbophyllinae Schltr. (tribe Dendrobieae Endl.) included the large genus *Bulbophyllum* along with smaller genera, such as *Cirrhopetalum* Lindl., *Drymoda* Lindl., *Pedilochilus* Schltr., *Sunipia* Buch.-Ham. ex Sm., and *Trias* Lindl. (Dressler, 1993; Garay et al., 1994; Szlachetko and Margonska, 2001). Recent revisions treat all genera within the subtribe Bulbophyllinae in a more broadly defined *Bulbophyllum* and recognise 97 sections within the genus (Pridgeon et al., 2014; Vermeulen et al., 2014). Molecular phylogenetic studies have largely focused on species from specific geographic regions such as Madagascar and the Mascarenes (Fischer et al., 2007; Gamisch et al., 2015), the Neotropics (Smidt et al., 2013, 2011), and Peninsular Malaysia (Hosseini et al., 2012) or on taxonomic groups such as the *Cirrhopetalum* alliance (Hu et al., 2020), and few have taken a global perspective (e.g., Gamisch and Comes, 2019). These studies revealed a strong biogeographic pattern within the genus with four main clades that include species largely confined or endemic within one broader geographical area: 1) continental Africa, 2) Madagascar and the Mascarene Islands, 3) Southern America, or 4) Asia (Fischer et al., 2007; Gamisch et al., 2015; Gamisch and Comes, 2019; Smidt et al., 2011). The Southern American clade, the Madagascan clade, and the continental African clade together form a highly supported lineage (Fischer et al., 2007; Gamisch et al., 2015; Gamisch and Comes, 2019; Smidt et al., 2013; 2011), in sister group position to the Asian clade (Fischer et al., 2007; Gamisch et al., 2015; Gamisch and Comes, 2019). Previous molecular phylogenetic studies have mainly elucidated relationships within Madagascan, continental African and Neotropical sections and within the *Cirrhopetalum* alliance (Fischer et al., 2007; Gamisch et al., 2015; Smidt et al., 2011; Hu et al., 2020). However, evolutionary relationships within the Asian clade, which also includes taxa from the Australasian and Pacific regions, are still poorly understood, and the monophyly of sections within this clade has remained largely untested within a phylogenetic framework.

The study of hyper diverse groups such as *Bulbophyllum* requires a robust phylogenetic framework to assess monophyly of its infrageneric taxa. High throughput sequencing approaches facilitate the establishment of such a framework phylogeny to clarify broader evolutionary relationships and to assess the monophyly of infrageneric taxa and their trait evolution (Hassemer et al., 2019; van Kleinwee et al., 2022, Nargar et al. 2022). However, phylogenomic studies which provide insights into broad-level evolutionary relationships within the Asian clade of *Bulbophyllum* are still lacking. This hampers progress in understanding diversification of its evolutionary lineages in time and space and trait evolution within this highly diverse genus.

Within *Bulbophyllum*, section *Adelopetalum* has a unique distribution, being the only section with a centre of diversity in Australia (Brummit, 2001; Pridgeon et al., 2014, Vermeulen, 1993), and thus presents an interesting study case for range evolution within *Bulbophyllum*. The section comprises twelve tropical to temperate epi-lithophytic species. Nine species occur along Australia’s east coast in the montane forest communities of the Great Dividing Range, with one species (*B. argyropus*) also found on Australian islands (Lord Howe Island, and Norfolk Island). Two species are endemic to the montane forests of New Caledonia (*B. corythium, B. lingulatum*) and one to the lowland coastal forests of New Zealand (*B. tuberculatum*). The section was circumscribed based on morphological affinities recognised among ten species from Australia and New Caledonia previously assigned to *Bulbophyllum* sections *Desmosanthes, Racemosae* and *Sestochilus* (Dockrill, 1969; 1992; Vermeulen, 1993). Subsequent treatments recognised two additional species within the section, *B. weinthalii* and *B. exiguum* (Jones and Clements, 2002; Clements and Jones, 2006). Section *Adelopetalum* is characterised by plants having thin creeping rhizomes adpressed to the host, anchored by filamentous roots with small pseudobulbs that are crowded to widely spaced, and a small single flat leaf arising from the apex of the pseudobulb. The inflorescence is single to few-flowered, with small white, cream or yellow flowers, sometimes with red or purple patterns. The petals are smaller than the sepals but similar in shape, with the bases of the lateral sepals fused to the column foot. The fleshly three-lobed labellum is firmly hinged to the apex of the column foot.

Previous cladistic analysis of sect. *Adelopetalum* based on morphological characters resolved two main clades within the section, differentiated by the size and shape of the lower margin of the stelidia: the filiform column appendages typical for most *Bulbophyllum* (Vermeulen, 1993). Previous molecular phylogenetic studies based on the nuclear ribosomal ITS region (ITS1 + 5.8S + ITS2) included two to three representatives of the section (Gamisch et al., 2015; Gamisch and Comes, 2019), placing these in an early diverging position within the Asian clade. However, phylogenetic placement of the section within *Bulbophyllum* was not strongly supported (PP<90, BS 97) and thus requires further study. Further, phylogenetic relationships within sect. *Adelopetalum* and its ancestral range evolution are poorly understood and have not yet been investigated using a molecular phylogenetic approach.

The aims of this study were to 1) build a phylogenomic framework for *Bulbophyllum* with focus on the Asian clade 2) assess the monophyly and phylogenetic placement of sect. *Adelopetalum* within *Bulbophyllum*; 3) to infer interspecific relationships within sect. *Adelopetalum* 4) and to reconstruct the range evolution of sect. *Adelopetalum*.

## 2 Materials and methods

### 2.1 Sampling

In total, 136 orchid samples representing 114 species were included in the study. From the Asian, Australasian and Pacific regions a broad sampling was included representing 41 sections, i.e. 60% of sections recognised from these regions in the most recent treatment of the group (Pridgeon et al., 2014). From the Australasian region, all *Bulbophyllum* species were sampled (Australia: 30, New Zealand: 2). For *Bulbophyllum* sect. *Adelopetalum*, 28 samples were included, representing all 12 species recognised for the section. The morphologically closely related sect. *Minutissima* was included with nine samples representing five species comprising all four Australasian and Pacific species and two (of ca. 19) tropical Asian species. Sampling of representative species for the Afrotropical and Neotropical clades of *Bulbophyllum* was informed by previous phylogenetic studies (Fischer et al., 2007; Smidt et al., 2011; Gamisch and Comes, 2019). Species names follow the accepted taxonomy based on the World Checklist of Selected Plant Families (WCSP, 2022) and sectional taxonomy IOSPE (2022). Exceptions were made for *B. exiguum* which was placed in section *Adelopetalum* and *B. wolfei* which was placed in section *Polymeres* based on Jones and Clements (2002) and Clements and Jones (2006). The outgroup comprised representatives of subtribe Dendrobineae which is sister to Bulbophyllinae, and tribes Malaxideae, Arethuseae, Nervilleae, and Neottieae based on previous molecular phylogenetic studies (Givnish et al., 2015; Górniak et al., 2010, Serna-Sánchez et al. 2021). Details of the plant material studied, voucher information and the number of loci included for each sample are provided in Supplementary Material S1 and a complete list of loci analysed is provided in Supplementary Material S2.

### 2.2 DNA extraction, amplification, and sequencing

Total genomic DNA was extracted from ca. 10 to 20 mg silica-dried leaf material. Extractions were carried out with commercial extraction kits (Qiagen DNeasy plant kit, Venlo, Netherlands; ChargeSwitch gDNA plant kit, Invitrogen, Carlsbad, USA) following the manufacturer’s protocols or using the CTAB method (Doyle and Doyle, 1990), with modifications as described in Weising et al. (2005). Sequence data was generated using both Sanger sequencing (46 samples) for the nuclear ribosomal ITS region (ITS1, 5.8s, ITS2) and two plastid genes (*mat*K, *ycf*1) and shotgun high-throughput sequencing (89 samples) to extract 70 plastid coding sequences (CDS) and the nuclear ribosomal DNA cistron (Supplementary Material S2) for subsequent analyses. Libraries for high-throughput sequencing were constructed from 50 to 100 ng total DNA using the TruSeq Nano DNA LT library preparation kit (Illumina, San Diego, USA) for an insert size of 350 base pairs (bp) and paired-end reads following the manufacturer’s protocol. Libraries were multiplexed 96 times and DNA sequencing with 125 bp paired-end reads was carried out on an Illumina HiSeq 2500 platform at the Australian Genomic Research Facility, Melbourne (Australia).

For Sanger sequencing, amplifications for ITS were carried out with primers 17F and 26SER (Sun et al., 1994), for *mat*K with the primers 19F and, 1326R (Cuénoud et al., 2002), and for *ycf1* with primers 3720F, intR, intF, 5500R (Neubig et al., 2008). PCR reaction protocols are provided in Supplementary Material S3. Sequencing reactions were carried out using the amplification primers and sequencing was conducted on an AB3730xl 96-capillary sequencer (Australian Genome Research Facility, Brisbane, Australia).

### 2.3 Assembly and alignment

Sequences were assembled and edited in Geneious R10 (Kearse et al., 2012). Illumina sequences were assembled to a reference set of plastid CDS extracted from *Dendrobium catenatum* (GenBank accession numbers KJ862886) and for *ycf*68 from *Anoectochilus roxburghii* (KP776980). To build a reference for the nuclear ribosomal ITS-ETS region, Illumina reads of *B. boonjee* (CNS_G07175) were first mapped to the ITS-ETS region of *Corallorhiza trifida* (JVF2676a). To extend the region assembled, the *B. boonjee* Illumina reads were then mapped to the *B. boonjee* consensus sequence generated in the initial step, yielding a *B. boonjee* reference of the nuclear ribosomal DNA cistron (5′ETS, 18s, ITS1, 5.8s, ITS2, 28s, 3′ETS). Illumina sequences for all other samples were assembled against the *B. boonjee* nuclear ribosomal DNA cistron reference. Assemblies were carried out with the highest quality threshold and a minimum coverage of ten reads. The quality of the assemblies was checked and edited manually where required. Sequences were deposited in GenBank and ENA. For Sanger sequences, bidirectional reads were assembled in Geneious and edited manually. Additional sequences were sourced from DRYAD (https://doi.org/10.5061/dryad.n9r58) for *Coelogyne flaccida* (Givnish et al., 2015). DNA sequences were aligned using MAFFT v.7.222 (Katoh et al., 2005, 2002) with the default settings, visually inspected and then concatenated into a nuclear and plastid supermatrix, respectively. The nuclear supermatrix included 136 accessions, partitioned into coding and non-coding regions (alignment length: 6,341bp, number of parsimony informative sites: 995 (16%)); and the plastid supermatrix included 130 accessions, and 70 plastid coding regions, partitioned by gene and codon position (alignment length: 61,553bp, number of parsimony informative sites: 5,789 (9%)). A plastid dataset comprised of high throughput sequencing data only (excluding Sanger sequences) was produced including 90 accessions, and 70 plastid coding regions (alignment length: 61,553bp, number of parsimony informative sites: 5682 (9%) and analysed separately.

For divergence time estimations, the plastid supermatrix was reduced to one representative per species (indicated by an asterisk in Supplementary Material S1), comprising 111 accessions (alignment length: 60,984 bp, number of parsimony informative sites: 5,755 (9%)).

### 2.4 Phylogenetic analysis

Phylogenetic relationships were inferred using maximum likelihood (ML) in IQ-TREE v. 1.6.12 (Nguyen et al., 2015). The best-fit partition scheme and nucleotide substitution model for each partition was determined with IQ-TREE’s ModelFinder (Kalyaanamoorthy et al. 2017) based on the Akaike information criterion (AIC) (Akaike, 1974). Nodal support was assessed based on 1000 replicates of ultrafast bootstrap approximation with clades receiving >95 ultrafast bootstrap support (UFBS) considered as well supported (Minh et al., 2013; Hoang et al., 2018).

### 2.5 Divergence time estimation

Divergence times were estimated based on the plastid dataset in Beast2 v. 2.4.8 (Bouckaert et al., 2014) applying the best fit partition scheme and substitution model as determined by IQ-TREE’s ModelFinder. We tested two molecular clock models: 1) strict clock (Zuckerkandl and Pauling, 1965) and 2) relaxed lognormal clock (Drummond et al., 2006) and two models of speciation and extinction: 1) Yule and 2) birth-death (Yule, 1925; Gernhard et al., 2008). Three secondary calibration points were used applying priors with a normal distribution and mean ages and 95% higher posterior density (HDP) intervals based on the results of a family-wide molecular clock analysis by Chomicki et al. (2015). The root age was set to 55.02 Ma (HDP: 42.0–68.0). The next secondary calibration point was applied to the last common ancestor of Dendrobineae, Malaxideae, and Arethuseae and was set to 47.77 Ma (HDP: 36.4–59.1). Monophyly was constrained for this node consistent with relationships reconstructed in previous phylogenetic analyses (Chomicki et al. 2015; Givnish et al., 2015). The last secondary calibration was set at the stem node of Dendrobieae and Malaxideae with 38.68 Ma (HDP: 30.8–46.6). An additional calibration based on the fossil *Dendrobium winikaphyllum* (Conran et al., 2009) was applied to the stem node of the Australasian *Dendrobium* clade (*D. macropus, D. cunninghamii*, and *D. muricatum*), using a uniform distribution with an infinite maximum age and the minimum age constrained to 20.4 Ma, based on the minimum age of the strata containing the fossil (Mildenhall et al. 2014). Ten independent Beast analyses were run for 30 million MCMC generations, with trees sampled every 3×10^4^ generations. To assess convergence of independent runs and determine burn-in fractions, log files were assessed in Tracer v.1.7.1 (Rambaut and Drummond, 2007). Log and trees files from independent runs were combined in LogCombiner (from the Beast package) with a cumulative burn-in fraction of 10%-31% and the sampling frequency set to generate at least 10,000 tree and log files (Drummond and Bouckaert, 2015). The combined log file was assessed in Tracer to ensure the effective sample size of all parameters was above 200. An additional five independent Beast runs were conducted for the final analysis using a relaxed log normal clock with birth death speciation to achieve an effective sample size above 200 for the ucldmean parameter. A maximum clade credibility tree was generated in TreeAnnotator (Beast package) with median node heights. To compare clock and speciation models, the Akaiki information criterion by MCMC app from the BEAST 2 package v 2.6.2 was used to measure the AICM for the combined MCMC runs generated in the BEAST analysis for each model (Supplementary Material S4).

### 2.6 Ancestral range analysis

Species distributions were extracted from WCSP (2022). Biogeographic areas were largely delineated based on botanical continents defined by Brummit (2001). The subcontinental regions of Papuasia, Australia and New Zealand were recognised to allow a more fine-scaled resolution of range evolution in section *Adelopetalum* (Brummit 2001). The following seven biogeographic areas were coded: a, Africa; b, temperate Asia; c, tropical Asia; d, Papuasia; e, Australia; f, New Zealand, and g, Pacific. Ancestral ranges were estimated in RASP v. 4.0 (Yu et al., 2015) with the BioGeoBEARS package (Matzke, 2013) based on the maximum clade credibility tree obtained from the Beast analysis of the plastid supermatrix, pruned of the outgroups to Dendrobieae. Three models of range evolution were tested: the dispersal-extinction cladogenesis model (DEC) (Ree and Smith, 2008), a ML version of Ronquist’s parsimony dispersal-vicariance (DIVA; Ronquist, 1997), termed DIVALIKE (Matzke, 2013), and a simplified likelihood interpretation of the Bayesian “BayArea” program (Landis et al., 2013) known as BAYAREALIKE (Matzke, 2013). No constraints were applied to dispersal direction and the maximum number of ranges was set to five based on the maximum number of observed areas in extant species. Likelihood values were compared and the model of best fit determined by AIC score (Akaike, 1974) was used to infer the marginal probabilities of alternative ancestral ranges at each node in the phylogeny (Supplementary Material S5).

## 3 Results

### 3.1 Phylogenetic relationships

#### 3.1.1 Phylogenetic relationships – Plastid data

The ML phylogeny inferred from the 70 loci plastid supermatrix provided strong support for the monophyly of *Bulbophyllum* and its sister group relationship to *Dendrobium* (Fig. 1). Section *Adelopetalum* and *Minutissima s*.*s*. formed a highly supported clade, here termed the Adelopetalum/Minutissima clade, which was resolved in sister group position to the remainder of the genus (ultrafast bootstrap support/UFBS 98) (Fig. 1, Fig. 2). Within the Adelopetalum/Minutissima clade, all *Adelopetalum* species plus *B. pygmaeum* (sect. *Minutissima*) formed a highly supported lineage (UFBS 100), here termed the Adelopetalum clade. Within the Adelopetalum clade several highly supported groups were resolved: 1) the argyropus clade consisting of *B. argyropus, B. corythium* and *B. tuberculatum* (UFBS 100), reconstructed in a highly supported sister group relationship to *B. weinthalii* (UFBS 100); 2) the bracteatum clade, including *B. boonjee, B. bracteatum*, and *B. elisae* (UFBS 99); and 3) the newportii clade comprised of *B. exiguum, B. lageniforme, B. lilianae, B. lingulatum*, and *B. newportii* (UFBS 100). Relationships among *B. pygmaeum*, the argyropus clade + *B. weinthalii*, bracteatum and newportii clades received weak support. Sister to the Adelopetalum clade was the highly supported Minutissimum clade comprised of three species of sect. *Minutissima* (*B. globuliforme, B. keekee, B. minutissimum*), including the type species of the section (UFBS 100) (Fig. 1). Section *Minutissima* was identified as polyphyletic, with sect. *Minutissima* species placed within the Adelopetalum clade (*B. pygmaeum*) and the Asian clade (*B. mucronatum, B. moniliforme*). Within the Asian clade, sections *Beccariana, Brachyantha, Brachypus, Cirrhopetaloides, Cirrhopetalum, Desmosanthes, Oxysepala, Polymeres* and *Sestochilos* were identified as polyphyletic or paraphyletic. Phylogenetic relationships described here based on the plastid supermatrix (Fig.1, Fig. 2) are supported by reconstructions based on the 70 gene plastid dataset (Supplementary Material S6).

**Figure 1.**
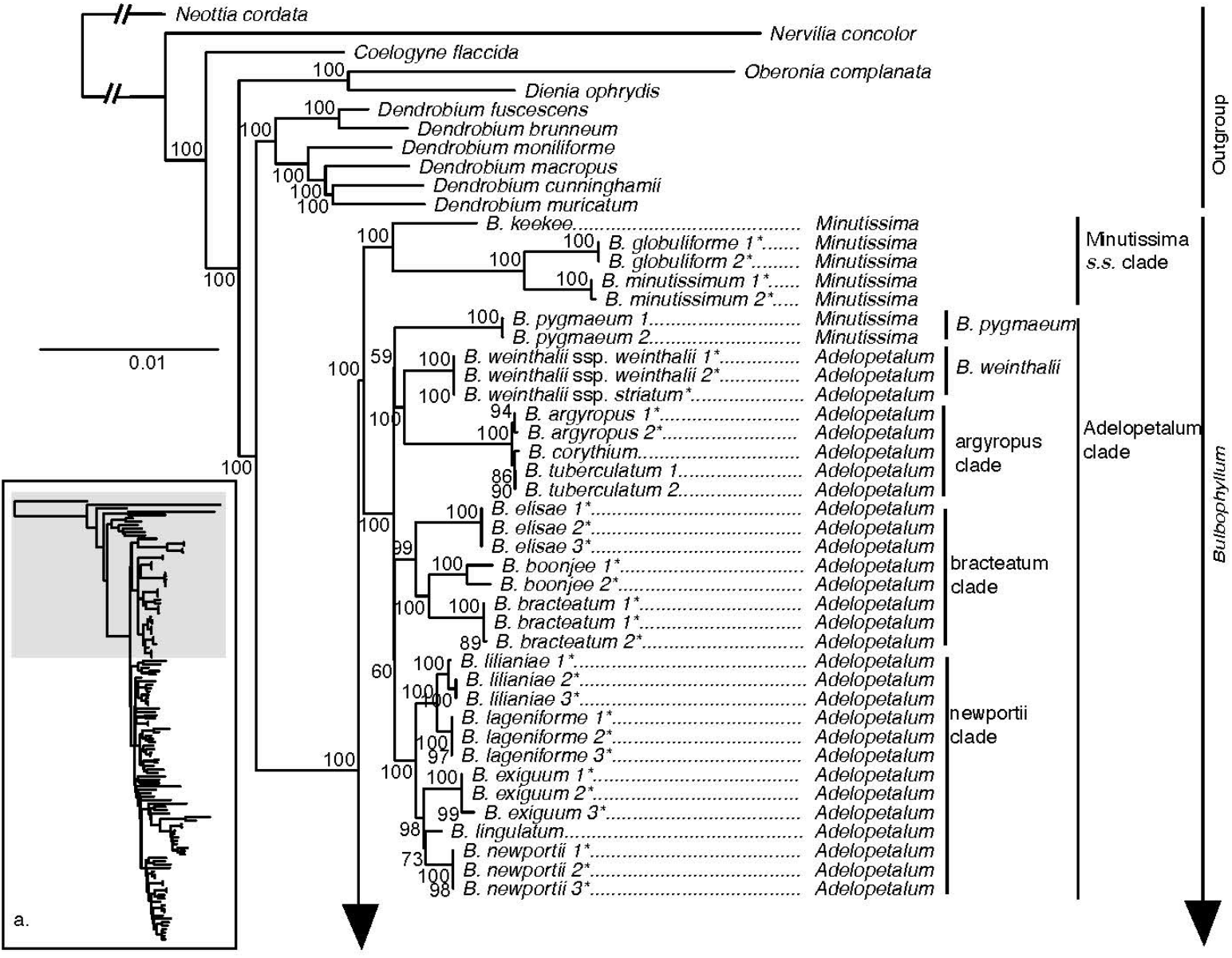
Maximum likelihood phylogenetic reconstruction of *Bulbophyllum* based on the supermatrix of 70 plastid coding regions in inset a. with *Bulbophyllum* sections *Adelopetalum* and *Minutissima* s.s. in detail. Ultrafast bootstrap values are given adjacent to nodes. Australian species are shown with an asterisk.

**Figure 2.**
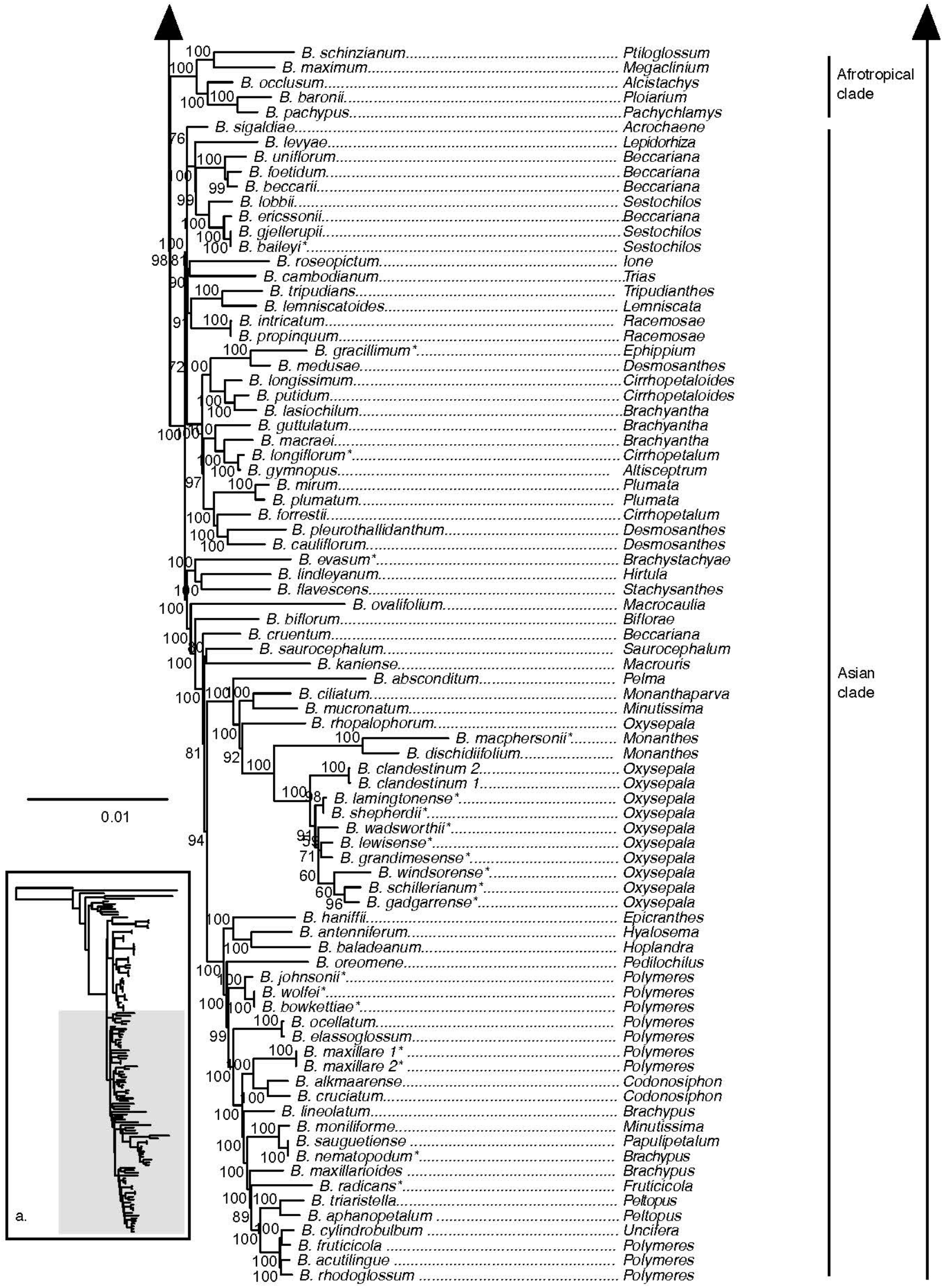
Maximum likelihood phylogenetic reconstruction of *Bulbophyllum* based on the supermatrix of 70 plastid coding regions in inset a. with Afrotropical and Asian *Bulbophyllum* clades in detail. Ultrafast bootstrap values are given adjacent to nodes. Australian species are shown with an asterisk.

Our analyses showed that sect. *Adelopetalum* does not share a close relationship with other Australasian *Bulbophyllum* species, such as those in sect. *Brachypus (B. nematopodum)*, sect. *Brachystachyae (B. evasum)*, sect. *Cirrhopetalum (B. longiflorum)*, sect. *Ephippium (B. gracillimum)*, sect. *Monanthes (B. macphersonii)*, sect. *Oxysepala (B. gadgarrense, B. grandimesense, B. lamingtonense, B. lewisense, B. schillerianum, B. shepherdii, B. wadsworthii, B. windsorense)*, sect. *Polymeres (B. bowkettiae, B. johnsonii, B. radicans, B. wolfei)*, and sect. *Sestochilus (B. baileyi)*. Australian species from each of these sections were placed in nine different positions within the Asian clade. Australian species from section *Polymeres* formed a highly supported clade (UFBS 100), while Australian species from sect. *Oxysepala* formed a moderately supported clade (UFBS 91) and together with the type species of section *Oxysepala* from Papuasia (*B. cladistinum*) formed a close relationship with the Australian representative of section *Monanthes* (*B. macphersonii*) (UFBS 100).

#### 3.1.2 Phylogenetic relationships – Nuclear data

The ML phylogeny based on the nuclear ribosomal DNA cistron was resolved with overall lower support compared to analyses based on 70 plastid loci supermatrix (Fig. 3, Fig. 4). Relationships among outgroup taxa were concordant with the plastid phylogeny and *Bulbophyllum* was resolved with maximum support. Within *Bulbophyllum*, the Afrotropical (UFBS 97), Asian (UFBS 100) and Adelopetalum/Minutissima (UFBS 100) clades were resolved with high to maximum support, however the relationships among them were poorly supported. Within the Adelopetalum/Minutissima clade, the highly supported clades revealed in the plastid phylogeny were also reconstructed based on the nuclear dataset (argyropus clade (UFBS 100), bracteatum clade (UFBS 93), minutissimum clade (UFBS 86), and newportii clade (UFBS 83), however relationships among these remained poorly supported. Similar to reconstructions based on the plastid phylogeny were the relationships among the argyropus, bracteatum and newportii clades, the poor support of *B. pygmaeum* and *B. weinthalii*, the polyphyly or paraphyly for sections *Beccariana, Brachyantha, Brachypus, Cirrhopetaloides, Cirrhopetalum, Desmosanthes, Minutissima, Oxysepala, Polymeres*, and *Sestochilos*; and that Australian species from sect. sect. *Brachypus*, sect. *Brachystachyae*, sect. *Cirrhopetalum*,, sect. *Ephippium*, sect. *Monanthes*, sect. *Oxysepala*,, sect. *Polymeres*, and sect. *Stenochilus* were placed in nine clades across the Asian clade.

**Figure 3.**
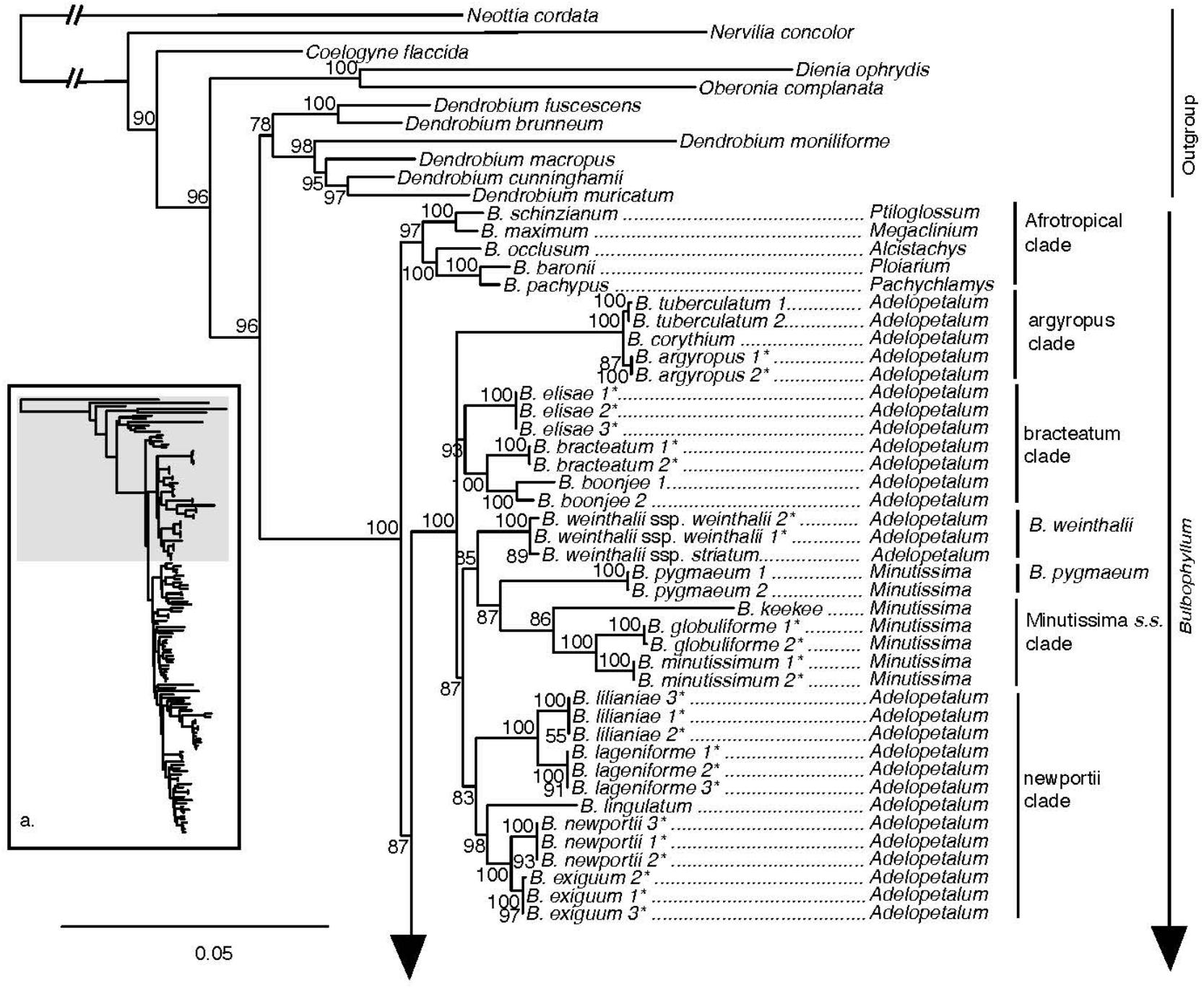
Maximum likelihood phylogenetic reconstruction of *Bulbophyllum* based on the nuclear ribosomal DNA cistron (5′ETS, 18s, ITS1, 5.8s, ITS2, 28s, 3′ETS) in inset a. with *Bulbophyllum* sections *Adelopetalum* and *Minutissima* in detail. Ultrafast bootstrap values are given adjacent to nodes. Australian species are shown with an asterisk.

**Figure 4.**
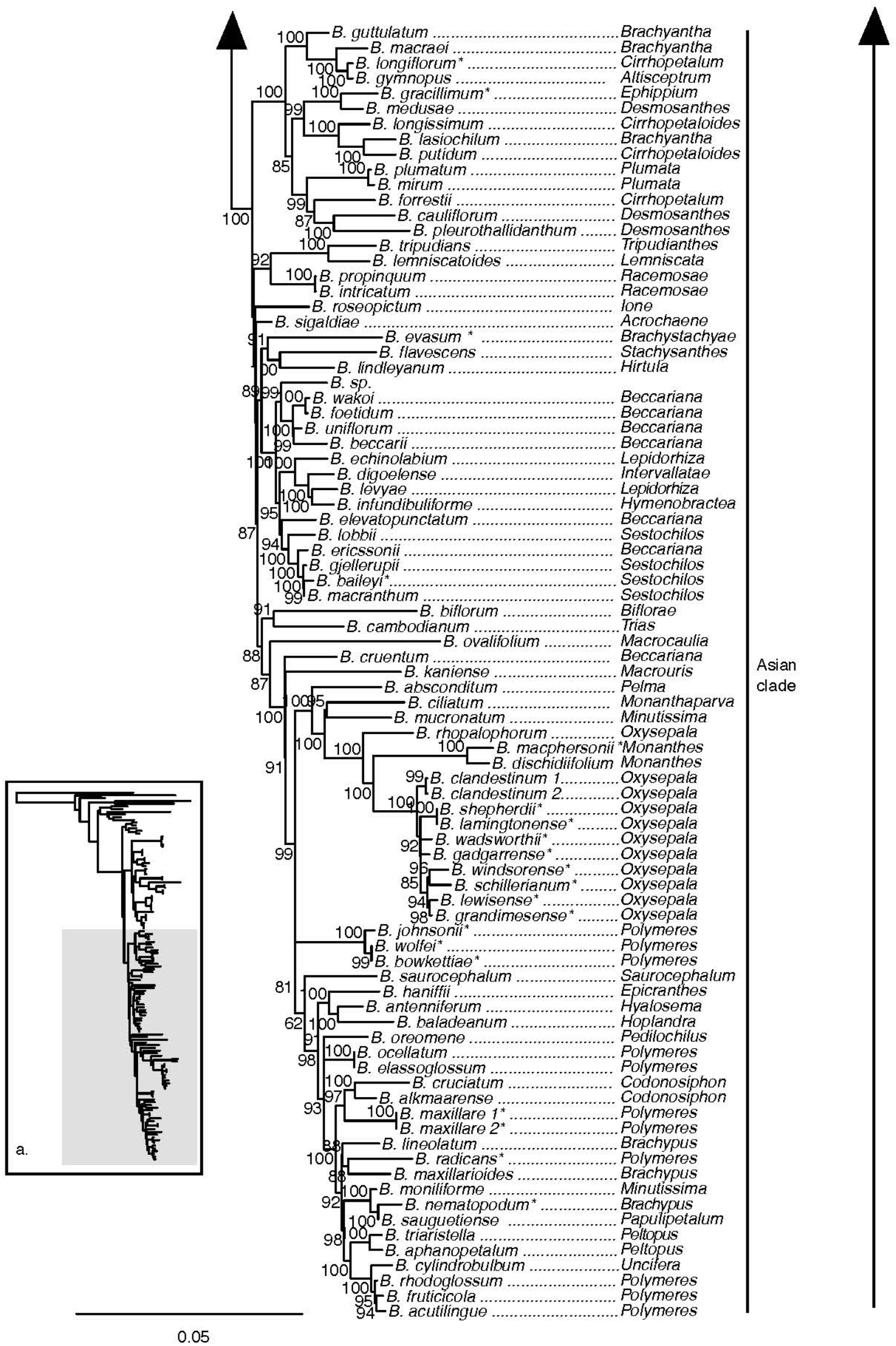
Maximum likelihood phylogenetic reconstruction of *Bulbophyllum* based on nuclear ribosomal DNA cistron (5′ETS, 18s, ITS1, 5.8s, ITS2, 28s, 3′ETS) in inset a. with Afrotropical and Asian *Bulbophyllum* clades in detail. Ultrafast bootstrap values are given adjacent to nodes. Australian species are shown with an asterisk.

### 3.2 Divergence time estimation

The divergence time analysis based on a relaxed log normal clock and birth death prior with speciation and extinction, which was identified as the model of best fit based on the Akaike information criterion (Supplementary Material S4), is presented here (Fig. 5) with the Asian and Afrotropical clades collapsed and the complete chronogram provided in Supplementary Material S7. The divergence time analysis based on the plastid dataset was well resolved and highly supported (Fig. 5, Supplementary Material S7). The divergence between *Bulbophyllum* and *Dendrobium* was estimated to have occurred during the early Oligocene, ca. 33.2 Ma (95% highest posterior probability density, HPD: 27.7–39.0). The crown of *Bulbophyllum*, constituting the divergence of the Adelopetalum/Minutissima clade from the remainder of the genus, was dated to the late Oligocene, ca. 24.9 Ma (HPD: 20.1–30.7). Divergence between the Asian clade and the Afrotropical clade was estimated to have taken place during the late Oligocene, ca. 24.3 Ma (HPD: 19.4–29.8) and diversification within the Asian clade was estimated from the mid Miocene 21.4 Ma (HPD: 17.2– 26.4 Ma) and the Afrotropical clade from 16.0 Ma (HPD: 10.4–21.7). The crown age of the Adelopetalum/Minutissima clade was dated to the late Oligocene, ca. 23.6 Ma (HPD: 18.6–29.1), with the split of the Minutissimum clade from the Adelopetalum clade. The crown age of the Adelopetalum clade was dated to the mid Miocene, ca. 15.3 Ma (HPD: 10.6–21.2). The stem branches of major lineages within the Adelopetalum clade were estimated to have diversified during the mid-Miocene: the bracteatum clade was dated to ca. 14.5 Ma (HPD: 10.0–20.4); the lineage giving rise to *B. weinthalii* to ca. 12.1Ma (6.9–17.6); the argyropus clade to ca. 12.1 Ma (6.9–17.6); and the newportii clade to ca. 14.9 Ma (HPD 10.3–20.7). Diversification among species within these lineages took place from the mid-Miocene onwards with the most recent divergence identified during the late Pleistocene among *B. argyropus, B. corythium*, and *B. tuberculatum*.

**Figure 5.**
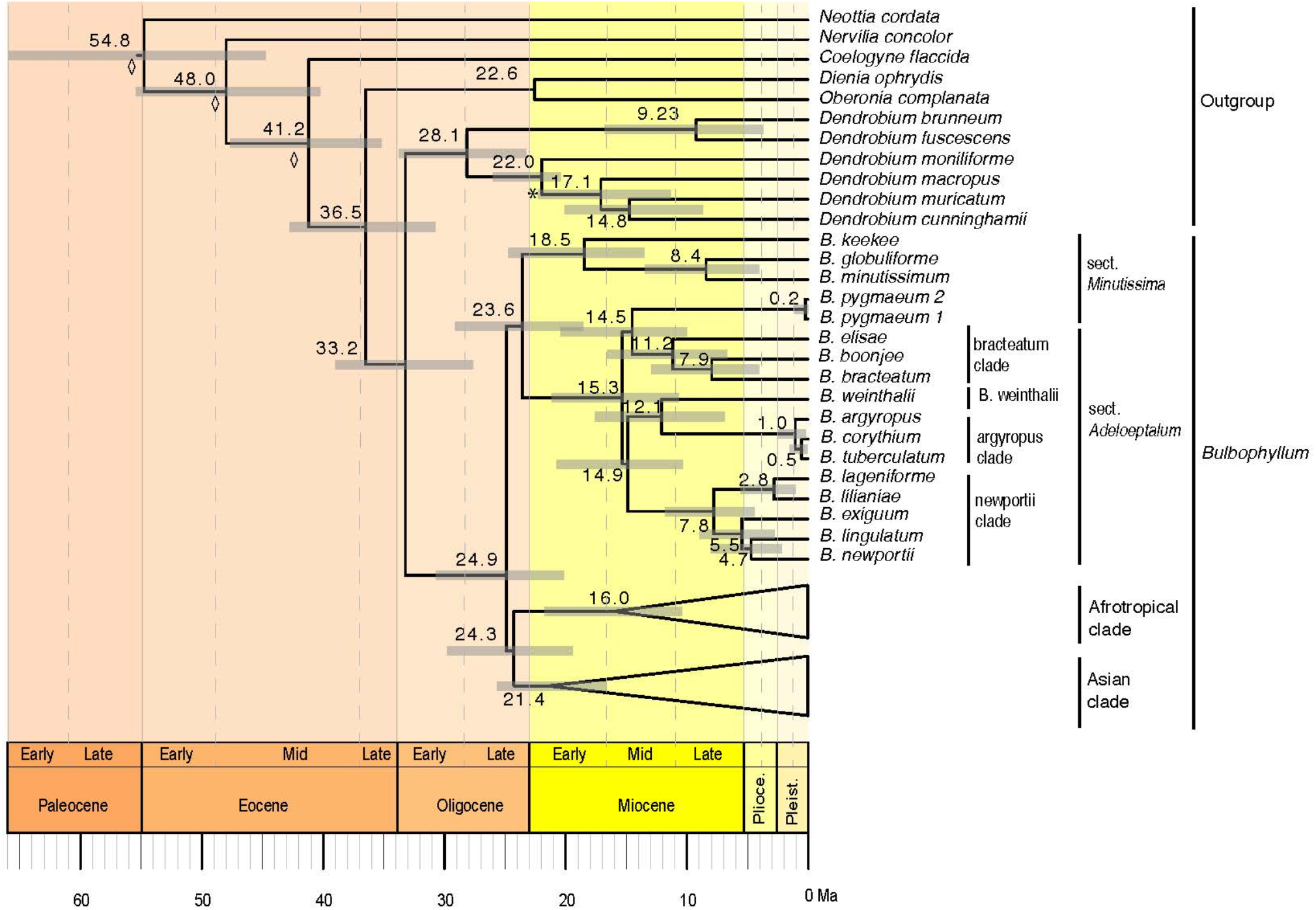
Maximum clade credibility chronogram for *Bulbophyllum* sect. *Adelopetalum* based on 70 plastid coding sequences, relaxed log normal clock and birth death prior. Divergence dates and 95% highest posterior density values are indicated adjacent to nodes. Grey bars indicate 95% highest posterior density. The asterisk denotes the node constrained with a fossil calibration point; the diamond shape denotes nodes which were constrained by secondary calibration points.

### 3.3 Ancestral range analysis

Model testing of the three biogeographic models (DEC, DIVALIKE, BAYAREALIKE) using the Akaike information criterion identified the BAYAREALIKE model as the model of best fit for the ancestral range estimation (Supplementary Material S5). Ancestral ranges estimated with the BAYAREALIKE model are presented here with the Asian and Afrotropical clades collapsed (Fig. 6). The complete chronogram is provided in Supplementary Material S8, marginal probabilities for ancestral ranges at all nodes in Supplementary Material S9 and node IDs in Supplementary Material S10.

**Figure 6.**
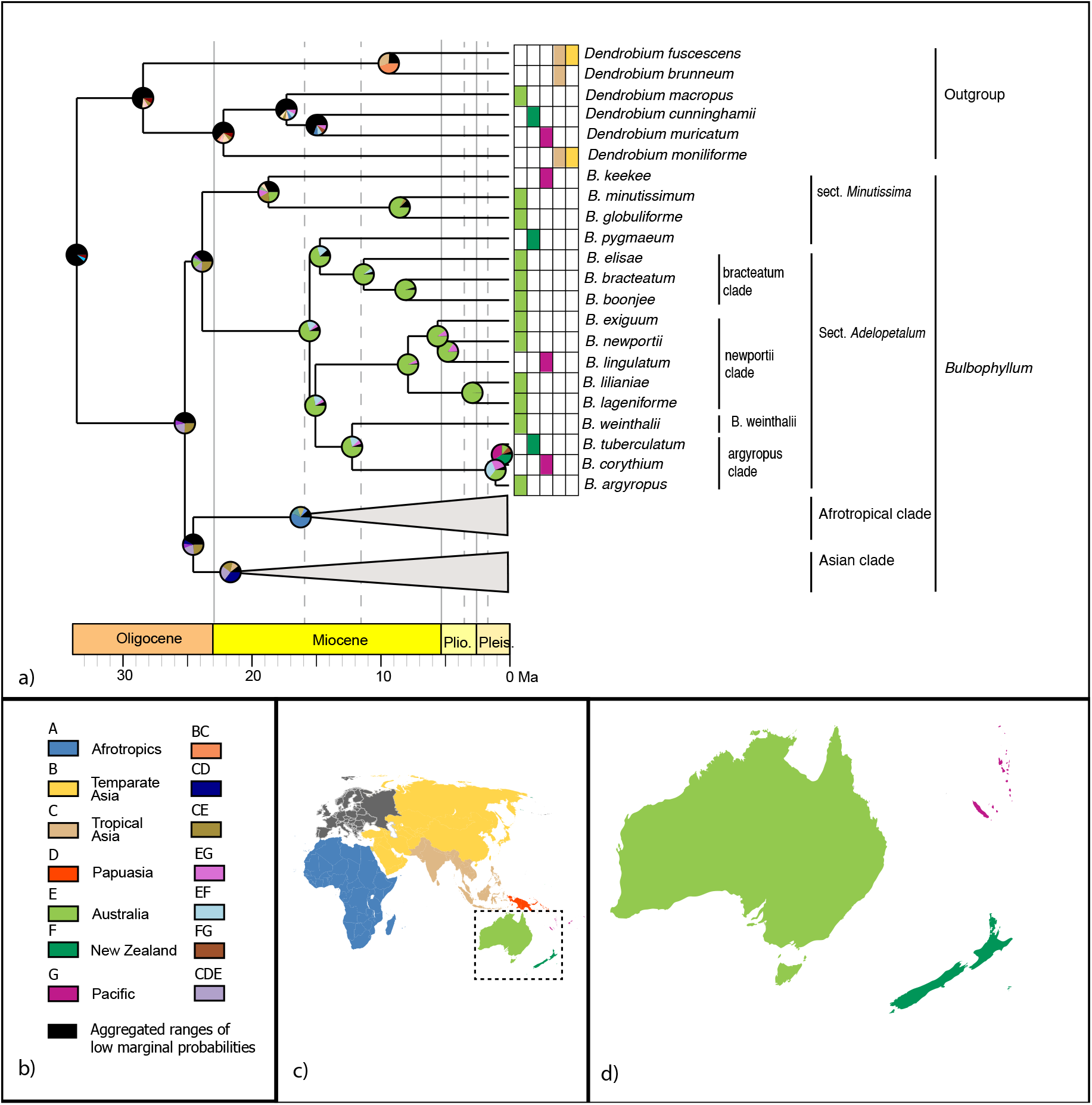
Range evolution of *Bulbophyllum* sect. *Adelopetalum*. a) ancestral area reconstruction based on a the BAYAREALIKE model with species extant distributions shown within the grid and pie charts at internal nodes representing marginal probabilities for alternative ancestral areas; b) legend of color-coded geographic regions and shared ancestral areas; c) world map of color-coded geographic regions delineated in the biogeographic analysis; d) detail of Australasian color-coded geographic regions.

Australia was reconstructed as the most likely ancestral range for the MRCA of the Adelopetalum clade (range probability (RP) 82) and all nodes within this lineage (RP 72-99) except for the argyropus clade (Fig. 6). Range shifts from Australia were inferred from the early Pliocene to the Pacific region (New Caledonia) in the newportii clade, in the lineage giving rise to *B. lingulatum*. Range shifts were also inferred from Australia to the Pacific region (New Caledonia) and New Zealand either in the lineage giving rise to the MRCA of the argyropus clade or subsequently within this lineage. Three alternative ancestral ranges were reconstructed for the MRCA of the argyropus clade: Australia (RP 36), or widespread distributions including Australia and New Zealand (RP 33) or Australia and New Caledonia (RP 26). Two alternative ranges were also reconstructed for the MRCA of *B. corythium* and *B. tuberculatum*: New Zealand (RP 41) and New Caledonia (RP 34). Considering these alternative scenarios, range shifts within the argyropus clade were estimated to have occurred sometime between the mid Miocene and late Pliocene (12.1–0.5 Ma). The ancestral range of MRCA of the Adelopetalum/Minutissima clade and Bulbophyllum remained unresolved in the ancestral range reconstruction. The most likely ancestral range for the MRCA of the Adeloptalum/Minutissima clade was a widespread distribution across Australia and tropical Asia (RP 26), while alternative ranges reconstructed included a widespread range including Australia, tropical Asia and Papuasia (RP 13) and Australia (RP 11). Two alternative ancestral ranges were reconstructed for the MRCA of Bulbophyllum, both widespread distributions including Australia and tropical Asia (RP 25.9) or Australia, tropical Asia and Papuasia (RP 21).

## 4 Discussion

### 4.1 Phylogenetic relationships

This study provided a broad plastid phylogenetic framework for Asian and Australasian sections of *Bulbophyllum* and revealed a close relationship between sections *Adelopetalum* and *Minutissima s*.*s*., that together form a highly supported early diverging lineage within the genus (Fig. 1, Fig. 2). Relationships based on 70 plastid genes support a sister group relationship between the *Adelopetalum*/*Minutissima* clade and the remainder of the genus (Asian + Afrotropical clades). Within the Adelopetalum/Minutissima clade, analyses based on our 70 plastid loci supermatrix showed a dichotomous split between the highly supported *Minutissima s*.*s*. and Adelopetalum clades. Species were reconstructed in each of these clades according to their sectional placement except for New Zealand endemic *B. pygmaeum* (sect. *Minutissima*), which was nested within the Adelopetalum clade, rendering section *Adelopetalum* paraphyletic. Section *Minutissima* was identified as polyphyletic, with the Australian (*B. minutissimum* (sect. type), *B. globuliforme*) and Pacific species (*B. keekee*) placed in the Minutissima clade and New Zealand species (*B. pygmaeum*) in the Adelopetalum clade while the Asian species, *B. mucronatum* and *B. moniliforme* were resolved within the Asian clade. Section *Minutissima* has undergone numerous taxonomic changes with treatments ranging from a narrower circumscription recognising species from Australia (Jones and Clements, 2001), to broader classifications including 23 species from Thailand, Indonesia, Australia, New Zealand, New Caledonia, and New Guinea (Pridgeon et al. 2014). Our phylogenetic analysis based on plastid and nuclear markers did not reconstruct a close relationship between sect. *Minutissima* species from the Australasian/Pacific region and Asian species *B. mucronatum* and *B. moniliforme*. Rather, in our analyses the Australasian species fell within the Adelopetalum/Minutissima clade while the Asian species were nested within the Asian clade. The results support morphological studies differentiating sect. *Minutissima* species from Australasia and Asia (Jones and Clements, 2001) and show minute pseudobulbs are a trait that has evolved more than once independently in the genus.

Within section *Adelopetalum*, phylogenetic analyses supported current species concepts, except for species within the argyropus clade, which exhibited shallow genetic differentiation (Fig. 1, Fig. 3). The species of the argyropus clade share morphological affinities and previous taxonomic treatments recognised up to three species within the group: *B. argyropus* (Australia’s east coast and off shore islands: Lord Howe Island, and Norfolk Island), *B. corythium* (New Caledonia), and *B. tuberculatum* (New Zealand) (Clements and Jones, 2002; Halle, 1981; Vermeulen, 1993). Divergence dating analyses shows this group represents a relatively recent radiation reconstructing divergence among species during the Pleistocene. Further studies are required to clarify species delimitation and dispersal patterns utilising population-level sampling and genomic techniques suited to resolving relationships among recently diverged lineages, such as reduced representation high-throughput approaches like ddRAD, DArT or target sequence capture methods (Peterson et al., 2012; Sansaloni et al., 2011; Weitemier et al., 2014; Folk et al., 2015; Bagley et al. 2020, Schmidt-Lebuhn et al., 2022).

Previous cladistic analysis based on morphological traits in sect. *Adelopetalum* found two main clades within the section, one comprising *B. lageniforme, B. lilianae, B. lingulatum*, and *B. newportii*, and the other uniting *B. argyropus, B. bracteatum*, and *B. elisae* (Vermeulen 1993). Phylogenetic relationships based on plastid and nuclear markers found strong to moderate support for the first clade recovered in the cladistic analysis, corresponding to the newportii clade in the present analyses (Fig 1, Fig. 3). The second group found in the cladistics analysis included three species placed in phylogenetic analyses within either the bracteatum clade (*<u>B. bracteatum, B. elisae</u>*) or the argyropus clade (*B. argyropus*). However, relationships among these two lineages remained unclear due to low support. The sister group relationship between *B. bracteatum* and *B. argyropus* recovered in the cladistic analysis was not supported in phylogenetic reconstructions based on molecular data, suggesting character states shared by these species may be homoplasious. Further studies using ancestral character reconstruction are required to test the phylogenetic utility of morphological traits utilised in prior studies.

While plastid phylogenomics has clarified major clades and intraspecific relationships within sect. *Adelopetalum* and broad-level relationships within *Bulbophyllum*, further studies are required. Non-monophyletic sections identified in the present study (e.g., sections *Beccariana, Brachyantha, Brachypus, Cirrhopetaloides, Cirrhopetalum, Desmosanthes, Minutissima* and *Polymeres*) (Fig. 2) and in previous molecular phylogenetic studies (Fischer et al., 2007; Smidt et al., 2011; Pridgeon et al. 2014; Hu 2020) highlight the need for further taxonomic revision within *Bulbophyllum*. Studies are required with an expanded sampling of the diverse Asian and Pacific taxa to increase our understanding of evolutionary relationships and assess sectional classification in more detail. Phylogenetic relationships reconstructed from the nuclear ribosomal DNA cistron were not strongly supported in line with previous molecular studies based in ITS (Gamisch et al., 2015; Gamisch and Comes, 2019, Hu et al., 2020). Approaches yielding higher number of nuclear markers such as target sequence capture, provide an opportunity to improve the understanding of evolutionary relationships in future studies. While assembling datasets with comprehensive species coverage within mega diverse groups such as *Bulbophyllum* remains a challenge, the present study provides an example of the use of a broad phylogenetic framework with targeted sampling within a section, to test the monophyly and phylogenetic placement of groups of interest.

### 4.2 Spatio-temporal evolution of *Bulbophyllum* sect. *Adelopetalum*

Our divergence time analysis and ancestral range estimations showed that *Bulbophyllum* sect. *Adelopetalum* represents an Australasian lineage that originated on the Australian continent during the late Oligocene to early Miocene (Fig. 5, Fig. 6). The Australian ancestral range is largely conserved within the lineage, indicating diversification among species has predominantly taken place on the Australian continent. The conservation of ancestral range observed within sect. *Adelopetalum* is consistent with previous phylogenetic analyses of *Bulbophyllum* that have shown a strong biogeographic signal among clades, being largely confined to biogeographic regions such as Madagascar, continental Africa and South America (Fischer et al., 2007; Gamisch et al., 2015; Gamisch and Comes, 2019; Smidt et al., 2011). The evolution of *Bulbophyllum* during the early Oligocene occurred subsequently to the breakup of Gondwana (Matthews, et al. 2016; Zahirovic et al. 2016), implicating long-distance dispersal (LDD) in the evolution of biogeographical lineages within the genus (Van den Berg, 2003, Smidt et al., 2011; Gamisch et al., 2015; Gamisch and Comes, 2019). Nevertheless, the conservation of ancestral ranges observed within the *Adelopetalum* lineage in this study and strong biogeographic signal among clades identified in previous studies indicate LDD with successful establishment and persistence has been relatively infrequent within *Bulbophyllum*. Although the minute wind-dispersed seeds of orchids have a high dispersal potential, successful establishment in a new area are limited by several factors, such as the presence of mycorrhizal partners necessary for germination and development, a suitable host or substrate and microclimatic conditions, and the availability of pollinators (Arditti and Ghani, 2000; Jersáková and Malinová, 2007; McCormick et al., 2012). Our results are consistent with previous studies that have identified in situ diversification as the dominant biogeographic process, despite evidence for LDD, and provide further support for the hypothesis that the complex requirements for successful establishment, rather than dispersal limitations, play an important role in constraining the geographic distribution of orchids (Givnish et al., 2016; Perez-Escobar and Chomicki et al. 2017).

Our phylogenetic analysis further resolved interspecific relationships in sect. *Adelopetalum* (Fig. 1). Divergence time estimation showed that divergence among species occurred mainly during the Miocene and Pliocene (Fig. 5), during a period of extensive changes to the distribution of forest vegetation on the Australian continent in response to drastic climatic changes. During the early Miocene, Australian vegetation diversified in response to aridification of the Australian continent and the abrupt shift to a cool dry climate during the mid-Miocene resulted in considerable fragmentation of rainforest habitats (Martin, 2006, Byrne et al. 2011). *Bulbophyllum* sect. *Adelopetalum* comprises epiphytic species that occur in mesic forest habitats and thus diversification and fragmentation of these habitats were likely drivers of allopatric lineage divergence within this group. Sister group relationships were identified between two species pairs with disjunct distributions in Australia’s northern wet tropical rainforests and south-eastern rainforests (*B. boonjee/B. bracteatum* and *B. newportii/B. exiguum*). These relationships support the hypothesis that the diversification and fragmentation of forest habitats in Australia has been an important driver of lineage divergence in Australia’s mesic biome (Byrne et al., 2011, Simpson et al., 2018).

Whilst the ancestral range was predominantly conserved within the *Adelopetalum* lineage, range expansion events were inferred from continental Australia across the Coral and Tasman Seas to New Caledonia in the lineage giving rise to *B. lingulatum*, and to New Zealand and New Caledonia in the argyropus clade (Fig. 6). New Caledonia and New Zealand each have a long history of isolation from Australia that predates the evolution of *Bulbophyllum*, indicating colonisation of these islands by *Bulbophyllum* species has been via LDD (Matthews et al., 2016). It remains unclear if LDD to New Zealand and New Caledonia in the argyropus clade occurred from the early Miocene in the lineage giving rise to the MRCA of the group or subsequently within this clade during the late Pleistocene, thus the spatio temporal evolution of this lineage requires further study.

The pattern of eastward dispersal observed in range shifts from Australia, across the Coral and Tasman Seas, is consistent with dispersal patterns inferred in other angiosperms, including *Abrotanella, Dendrobium, Dracophyllum, Hebe, Korthalsella, Leucopogon, Northofagus, Oreobolus, Pterostylis, Rytidosperma* (Chacón et al., 2006; Lavarack et al., 2000; Linder, 1999; Molvray et al., 1999; Swenson et al., 2001; Puente-Lelièvre et al., 2013; Wagstaff et al., 2010, 2006, 2002, Nargar et al. 2022). The bias towards eastward dispersal observed within section *Adelopetalum* among other plant groups may be facilitated by the predominant westerly winds occurring in the southern hemisphere that initiated after the rifting of Australia and South America from Antarctica during the Eocene (Sanmartín et al., 2007).

### 4.3 Conclusions

This study provided an important phylogenomic framework for the mega genus *Bulbophyllum* facilitating studies into trait and range evolution within the genus. Several Asian sections were resolved as paraphyletic warranting taxonomic revisions. Our plastid phylogenomic analyses revealed an early-diverging lineage within *Bulbophyllum*, composed of sect. *Adelopetalum* and sect. Minutissima *s*.*s*.. For *Bulbophyllum* sect. *Adelopetalum*, this study reconstructed an origin in the early Oligocene and identified the Australian continent as ancestral range. Species diversification within the section occurred predominantly on the Australian continent with fragmentation of mesic habitats during the Miocene identified as likely drivers of allopatric lineage divergence. Multiple independent long distance dispersal events were inferred from the Australian continent eastward to the islands of New Zealand and New Caledonia.

## 6 Author Contributions

Conceptualization: LS, KN, MAC; Data curation: LS, MAC; Formal Analysis: LS, HKO; Funding acquisition: LS, KN, DMC, MAC; Investigation and Methodology: LS; Supervision: KN, DCM, MAC; Writing – original draft: LS; Writing – review and editing: KN, HKO, DCM, MAC. All authors approved of the final version of the manuscript.

## 8 Acknowledgments

The authors wish to acknowledge the use of the Next-Generation Sequencing services and facilities of the Australian Genomic Research Facilities (AGRF). Plant material was collected under kind permission of the KuKu Nyungkal traditional owners, Queensland Parks and Wildlife (permit numbers: WISP11258812, WITK11258712. We thank B. Gray, D.L. Jones, G. McCraith, M.T. Mathieson, B.P. Molloy, L. Roberts, and P.D. Ziesing Michael Harrison for contributing plant material to the study. We thank Guillaume Chomicki for providing files utilised for secondary calibration of our divergence dating analyses.

## References

Akaike, H., 1974. A new look at the statistical model identification. IEEE Trans. Automat. Contr. 19, 716–723. doi:10.1109/TAC.1974.1100705

Arditti, J., Ghani, A.K.A., 2000. Numerical and physical properties of orchid seeds and their biological implications. New Phytol. 145, 367–421. doi:10.1046/j.1469-8137.2000.00587.x

Bagley, J. C., Uribe-Convers, S., Carlsen, M. M., & Muchhala, N. (2020). Utility of targeted sequence capture for phylogenomics in rapid, recent angiosperm radiations: Neotropical Burmeistera bellflowers as a case study. Molecular Phylogenetics and Evolution, 152, 106769.

Bouckaert, R., Heled, J., Kühnert, D., Vaughan, T., Wu, C.-H., Xie, D., Suchard, M.A., Rambaut, A., Drummond, A.J., 2014. BEAST 2: A software platform for Bayesian evolutionary analysis. PLoS Comput. Biol. 10, e1003537. doi:10.1371/journal.pcbi.1003537

Brummitt, R.K., 2001. World geographical scheme for recording plant distributions, edition 2. Hunt Institute for Botanical Documentation, Carnegie Mellon University, Pittsburgh.

Byrne, M., Steane, D.A., Joseph, L., Yeates, D.K., Jordan, G.J., Crayn, D., Aplin, K., Cantrill, D.J., Cook, L.G., Crisp, M.D., Keogh, J.S., Melville, J., Moritz, C., Porch, N., Sniderman, J.M.K., Sunnucks, P., 2011. Decline of a biome : evolution, contraction, fragmentation, extinction and invasion of the Australian mesic zone biota. J. Biogeogr. 38, 1–22. doi:10.1111/j.1365-2699.2011.02535.x

Chacón, J., Madriñán, S., Chase, M.W., Bruhl, J.J., 2006. Molecular phylogenetics of Oreobolus (Cyperaceae) and the origin and diversification of the American species. Taxon 55, 359–366. doi:10.2307/25065583

Chomicki, G., Bidel, L.P.R., Ming, F., Coiro, M., Zhang, X., Wang, Y., Baissac, Y., Jay-Allemand, C., Renner, S.S., 2015. The velamen protects photosynthetic orchid roots against UV-B damage, and a large dated phylogeny implies multiple gains and losses of this function during the Cenozoic. New Phytol. 205, 1330–1341. doi:10.1111/nph.13106

Conran, J.G., Bannister, J.M., Lee, D.E., 2009. Earliest orchid macrofossils: early Miocene Dendrobium and Earina (Orchidaceae: Epidendroideae) from New Zealand. Am. J. Bot. 96, 466–74. doi:10.3732/ajb.0800269

Cuénoud, P., Savolainen, V., Chatrou, L.W., Powell, M., Grayer, R.J., Chase, M.W., 2002. Molecular phylogenetics of Caryophyllales based on nuclear 18S rDNA and plastid rbcL, atpB, and matK DNA sequences. Am. J. Bot. 89, 132–44. doi:10.3732/ajb.89.1.132

Dockrill, A. W. (1969, 1992). Australian Indigenous Orchids: The epiphytes, the tropical terrestrial species. Surrey Beatty & Sons.

Doyle, J.J., Doyle, J.L., 1990. Isolation of plant DNA from fresh tissue. Focus 12, 13–15.

Dressler, R.L., 1993. Phylogeny and classification of the orchid family. Cambridge University Press.

Drummond, A. J., & Bouckaert, R. R. (2015). Bayesian evolutionary analysis with BEAST. Cambridge University Press.

Drummond, A.J., Ho, S.Y.W., Phillips, M.J., Rambaut, A., Rambaut, A., 2006. Relaxed Phylogenetics and dating with confidence. PLoS Biol. 4, e88. doi:10.1371/journal.pbio.0040088

Fischer, G. a, Gravendeel, B., Sieder, A., Andriantiana, J., Heiselmayer, P., Cribb, P.J., Smidt, E.D.C., Samuel, R., Kiehn, M., 2007. Evolution of resupination in Malagasy species of Bulbophyllum (Orchidaceae). Mol. Phylogenet. Evol. 45, 358–76. doi:10.1016/j.ympev.2007.06.023

Folk, R. A., Mandel, J. R., & Freudenstein, J. V. (2015). A protocol for targeted enrichment of intron-containing sequence markers for recent radiations: A phylogenomic example from Heuchera (Saxifragaceae). Applications in Plant Sciences, 3(8), 1500039.

Frodin, D.G., 2004. History and concepts of big plant genera. Taxon 53, 753–756. doi:10.2307/4135449

Gamisch, A., Fischer, G.A., Comes, H.P., 2015. Multiple independent origins of auto-pollination in tropical orchids (Bulbophyllum) in light of the hypothesis of selfing as an evolutionary dead end. BMC Evol. Biol. 15, 192–209. doi:10.1186/s12862-015-0471-5

Gamisch, A., Comes, H.P., 2019. Clade-age-dependent diversification under high species turnover shapes species richness disparities among tropical rainforest lineages of Bulbophyllum (Orchidaceae). BMC Evol. Biol. 19, 93. doi:10.1186/s12862-019-1416-1

Garay, L.A., Hamer, F., Siegerist, E.S., 1994. The genus Cirrhopetalum and the genera of the Bulbophyllum alliance. Nord. J. Bot. 14, 609–646. doi:10.1111/j.1756-1051.1994.tb01080.x

Gernhard, T., 2008. The conditioned reconstructed process. J. Theor. Biol. 253, 769–778. doi:10.1016/j.jtbi.2008.04.005

Givnish, T.J., Spalink, D., Ames, M., Lyon, S.P., Hunter, S.J., Zuluaga, A., Doucette, A., Caro, G.G., McDaniel, J., Clements, M.A., Arroyo, M.T.K., Endara, L., Kriebel, R., Williams, N.H., Cameron, K.M., 2016. Orchid historical biogeography, diversification, Antarctica and the paradox of orchid dispersal. J. Biogeogr. 43, 1905–1916. doi:10.1111/jbi.12854

Givnish, T.J., Spalink, D., Ames, M., Lyon, S.P., Hunter, S.J., Zuluaga, A., Iles, W.J.D., Clements, M.A., Arroyo, M.T.K., Leebens-Mack, J., Endara, L., Kriebel, R., Neubig, K.M., Whitten, W.M., Williams, N.H., Cameron, K.M., 2015. Orchid phylogenomics and multiple drivers of their extraordinary diversification. Proc. R. Soc. B Biol. Sci. 282, 20151553–20151562. doi:10.1098/rspb.2015.1553

Górniak, M., Paun, O., Chase, M.W., 2010. Phylogenetic relationships within Orchidaceae based on a low-copy nuclear coding gene, Xdh: Congruence with organellar and nuclear ribosomal DNA results. Mol. Phylogenet. Evol. 56, 784–95. doi:10.1016/j.ympev.2010.03.003

Halle, N., 1981. New data and carpograms of orchids from New Caledonia. Adansonia 20, 355–368.

Hassemer, G., Bruun-Lund, S., Shipunov, A. B., Briggs, B. G., Meudt, H. M., & Rønsted, N. (2019). The application of high-throughput sequencing for taxonomy: the case of Plantago subg. Plantago (Plantaginaceae). Molecular phylogenetics and evolution, 138, 156-173. doi.org/10.1016/j.ympev.2019.05.013

Hoang, D.T., Chernomor, O., von Haeseler A., Minh B.Q., and Vinh L.S., 2018. UFBoot2: Improving the ultrafast bootstrap approximation. Mol. Biol. Evol. 35, 518–522. https://doi.org/10.1093/molbev/msx281

Hosseini, S., Go, R., Dadkhah, K., & Nuruddin, A. A. (2012). Studies on maturase K sequences and systematic classification of Bulbophyllum in Peninsular Malaysia. Pak. J. Bot, 44(6), 2047–2054.

Hu, A. Q., Gale, S. W., Liu, Z. J., Suddee, S., Hsu, T. C., Fischer, G. A., & Saunders, R. M. (2020). Molecular phylogenetics and floral evolution of the Cirrhopetalum alliance (Bulbophyllum, Orchidaceae): Evolutionary transitions and phylogenetic signal variation. Molecular phylogenetics and evolution, 143, 106689.

IOSPE (2022, 6th July) Internet Orchid Species Photo Encyclopedia. http://www.orchidspecies.com/

Jersáková, J. and Malinová, T., 2007, Spatial aspects of seed dispersal and seedling recruitment in orchids.. New Phytol., 176, 237–241. doi:10.1111/j.1469-8137.2007.02223.x

Jones, D.L., Clements, M.A., 2001. Oncophyllum, a new genus of Orchidaceae from Australia. The Orchadian 13, 420–425.

Jones, D.L., Clements, M.A., 2002. Nomenclatural changes in the Australian and New Zealand Bulbophyllinae and Eriinae (Orchidaceae). The Orchadian 13, 498–501

Kalyaanamoorthy S., Minh B.Q., Wong T.K.F., von Haeseler A., and Jermiin L.S., 2017. ModelFinder: fast model selection for accurate phylogenetic estimates. Nat. Methods. 14, 587–589. DOI:10.1038/nmeth.4285

Katoh, K., Kuma, K., Toh, H., Miyata, T., 2005. MAFFT version 5: improvement in accuracy of multiple sequence alignment. Nucleic Acids Res. 33, 511–518. doi:10.1093/nar/gki198

Katoh, K., Misawa, K., Kuma, K., Miyata, T., 2002. MAFFT: a novel method for rapid multiple sequence alignment based on fast Fourier transform. Nucleic Acids Res. 30, 3059–66.

Kearse, M., Moir, R., Wilson, A., Stones-Havas, S., Cheung, M., Sturrock, S., Buxton, S., Cooper, A., Markowitz, S., Duran, C., Thierer, T., Ashton, B., Meintjes, P., Drummond, A., 2012. Geneious Basic: An integrated and extendable desktop software platform for the organization and analysis of sequence data. Bioinformatics 28, 1647–1649. doi:10.1093/bioinformatics/bts199

Landis, M. J., Matzke, N. J., Moore, B. R., & Huelsenbeck, J. P. (2013). Bayesian analysis of biogeography when the number of areas is large. Systematic biology, 62(6), 789–804.

Lavarack, B., Harris, W., Stocker, G., 2000. Dendrobium and its relatives. Timber Press, Portland.

Linder, H.P., 1999. Rytidosperma vickeryae — a new danthonioid grass from Kosciuszko (New South Wales, Australia): Morphology, phylogeny and biogeography. Aust. Syst. Bot. 12, 743–755. doi:10.1071/SB97046

Martin, H.A., 2006. Cenozoic climatic change and the development of the arid vegetation in Australia. J. Arid Environ. 66, 533–563. doi:10.1016/J.JARIDENV.2006.01.009

Matthews, K.J., Maloney, K.T., Zahirovic, S., Williams, S.E., Seton, M., Müller, R.D., 2016. Global plate boundary evolution and kinematics since the late Paleozoic. Glob. Planet. Change 146, 226–250. doi:10.1016/J.GLOPLACHA.2016.10.002

Matzke, N.J., 2013. BioGeoBEARS: Biogeography with Bayesian (and Likelihood) evolutionary analysis in R Scripts.

McCormick, M.K., Taylor, D., Juhaszova, K., Burnett, R.K., Whigham, D.F., O’neill, J.P., 2012. Limitations on orchid recruitment: not a simple picture. Mol. Ecol. 21, 1511–1523. doi:10.1111/j.1365-294X.2012.05468.x

Mildenhall, D.C., Kennedy, E.M., Lee, D.E., Kaulfuss, U., Bannister, J.M., Fox, B. and Conran, J.G., 2014. Palynology of the early Miocene Foulden Maar, Otago, New Zealand: Diversity following destruction. Review of Palaeobotany and Palynology, 204, pp.27–42.

Minh, B. Q., Nguyen, M. A. T., & von Haeseler, A. (2013). Ultrafast approximation for phylogenetic bootstrap. Molecular biology and evolution, 30(5), 1188–1195.

Molvray, M., Kores, P.J., Chase, M.W., 1999. Phylogenetic relationships within Korthalsella (Viscaceae) based on nuclear ITS and plastid trnL-F sequence data. Am. J. Bot. 86, 249–260. doi:10.2307/2656940

Nargar, K., O’Hara, K., Mertin, A., Bent, S., Nauheimer, L., Simpson, L., … & Clements, M. A. (2022). Evolutionary relationships and range evolution of greenhood orchids (subtribe Pterostylidinae): insights from plastid phylogenomics. Front. Plant Sci. 14, 1063174. doi.org/10.3389/fpls.2022.912089

Neubig, K.M., Whitten, A.W.M., Carlsward, B.S., Blanco, M.A., Endara, L., Williams, N.H., Moore, M., 2008. Phylogenetic utility of ycf1 in orchids: a plastid gene more variable than matK. Plant Syst. Evol. 277, 75–84. doi:10.1007/s00606-008-0105-0

Nguyen, L. T., Schmidt, H. A., Von Haeseler, A., & Minh, B. Q. (2015). IQ-TREE: a fast and effective stochastic algorithm for estimating maximum-likelihood phylogenies. Molecular biology and evolution, 32(1), 268-274. doi.org/10.1093/molbev/msu300

Peterson, B. K., Weber, J. N., Kay, E. H., Fisher, H. S., & Hoekstra, H. E. (2012). Double digest RADseq: an inexpensive method for de novo SNP discovery and genotyping in model and non-model species. PloS one, 7(5), e37135.

Pridgeon, A.M., Cribb, P.J., Chase, M.W., Rasmussen, F.N., 2014. Genera Orchidacearum Volume 6 Epidendroideae. Oxford University Press, New York, USA.

Puente-Lelièvre, C., Harrington, M.G., Brown, E.A., Kuzmina, M., Crayn, D.M., 2013. Cenozoic extinction and recolonization in the New Zealand flora: The case of the fleshy-fruited epacrids (Styphelieae, Styphelioideae, Ericaceae). Mol. Phylogenet. Evol. 66, 203–214. doi:10.1016/J.YMPEV.2012.09.027

Drummond, A. J., & Rambaut, A. (2007). BEAST: Bayesian evolutionary analysis by sampling trees. BMC evolutionary biology, 7(1), 1–8.

Rambaut, A., Drummond, A.J., 2007. Tracer v.1.5. Available from: http://beast.bio.ed.ac.uk/Tracer

Ree, R.H., Smith, S.A., 2008. Maximum likelihood inference of geographic range evolution by dispersal, local extinction, and cladogenesis. Syst. Biol. 57, 4–14. doi:10.1080/10635150701883881

Ronquist, F., 1997. Dispersal-vicariance analysis: a new approach to the quantification of historical biogeography. Syst. Biol 46, 195–203.

Ronquist, F., 2001. DIVA version 1.2. Computer program for MacOS and Win32.

Sanmartín, I., Wanntorp, L., Winkworth, R.C., 2007. West Wind Drift revisited: testing for directional dispersal in the Southern Hemisphere using event-based tree fitting. J. Biogeogr. 34, 398–416. doi:10.1111/j.1365-2699.2006.01655.x

Sansaloni, C., Petroli, C., Jaccoud, D., Carling, J., Detering, F., Grattapaglia, D., & Kilian, A. (2011). Diversity Arrays Technology (DArT) and next-generation sequencing combined: genome-wide, high throughput, highly informative genotyping for molecular breeding of Eucalyptus. In BMC proceedings (Vol. 5, No. 7, pp. 1-2). BioMed Central.

Serna-Sánchez, M. A., Pérez-Escobar, O. A., Bogarín, D., Torres-Jimenez, M. F., Alvarez-Yela, A. C., Arcila-Galvis, J. E., … & Arias, T. (2021). Plastid phylogenomics resolves ambiguous relationships within the orchid family and provides a solid timeframe for biogeography and macroevolution. Scientific reports, 11(1), 1–11.

Schmidt-Lebuhn, A. N., Egli, D., Grealy, A., Nicholls, J. A., Zwick, A., Dymock, J. J., & Gooden, B. (2022). Genetic data confirm the presence of Senecio madagascariensis in New Zealand. New Zealand Journal of Botany, 1–13.

Simpson, L., Clements, M.A., Crayn, D.M., Nargar, K., 2018. Evolution in Australia’s mesic biome under past and future climates: insights from a phylogenetic study of the Australian Rock Orchids (Dendrobium speciosum complex, Orchidaceae). Mol. Phylogenet. Evol. 118, 32–46. doi:10.1016/J.YMPEV.2017.09.004

Smidt, E.C., Gallo, L.W., Scatena, V.L., 2013. Leaf anatomical and molecular studies in Bulbophyllum section Micranthae (Orchidaceae) and their implications for systematics. Brazilian J. Bot. 36, 75–82. doi:10.1007/s40415-013-0008-3

Smidt, E.C., Borba, E.L., Gravendeel, B., Fischer, G.A., Berg, C. Van Den, 2011. Molecular phylogeny of the Neotropical sections of Bulbophyllum (Orchidaceae) using nuclear and plastid spacers. Taxon 60, 1050–1064.

Sun, Y., Skinner, D.Z., Liang, G.H., Hulbert, S.H., 1994. Phylogenetic analysis of Sorghum and related taxa using internal transcribed spacers of nuclear ribosomal DNA. Theor. Appl. Genet. 89, 26–32. doi:10.1007/BF00226978

Swenson, U., Backlund, A., McLoughlin, S., Hill, R.S., 2001. Nothofagus biogeography revisited with special emphasis on the enigmatic distribution of subgenus Brassospora in New Caledonia. Cladistics 17, 28–47. doi:10.1111/j.1096-0031.2001.tb00109.x

Szlachetko, D.L., Margonska, H.B., 2001. Genera et species Orchidalium. 3. Polish Bot. J. 461, 113–121.

Pérez-Escobar, O.A., Chomicki, G., Condamine, F.L., Karremans, A.P., Bogarín, D., Matzke, N.J., Silvestro, D., Antonelli, A., 2017. Recent origin and rapid speciation of Neotropical orchids in the world’s richest plant biodiversity hotspot. New Phytol. 215, 891–905. doi:10.1111/nph.14629

Van den Berg, C. 2003. Estudos em sistemática molecular na família Orchidaceae. Thesis presented for obtaining the title of Full Pro-fessor, Universidade Estadual de Feira de Santana, Bahia, Brazil.

van Kleinwee, I., Larridon, I., Shah, T., Bauters, K., Asselman, P., Goetghebeur, P., … & Veltjen, E. (2022). Plastid phylogenomics of the Sansevieria Clade of Dracaena (Asparagaceae) resolves a recent radiation. Molecular Phylogenetics and Evolution, 107404. doi.org/10.1016/j.ympev.2022.107404

Vermeulen, J.J., 1993. A taxonomic revision of Bulbophyllum sections Adelopetalum, Lepanthanthe, Macrouris, Pelma, Peltopus, and Unicifera (Orchidaceae). Orchid Monogr. 7.

Vermeulen, J.J., Schuiteman, A., De Vogel, E.F., 2014. Nomenclatural changes in Bulbophyllum (Orchidaceae; Epidendroideae). Phytotaxa 166, 101–113. doi:10.11646/phytotaxa.166.2.1

Wagstaff, S.J., Bayly, M.J., Garnock-Jones, P.J., Garnock-Jones, P.J., Albach, D.C., 2002. Classification, origin, and diversification of the New Zealand Hebes (Scrophulariaceae). Ann. Missouri Bot. Gard. 89, 38–63.

Wagstaff, S.J., Breitwieser, I., Swenson, U., 2006. Origin and relationships of the Austral genus Abrotanella (Asteraceae) inferred from DNA sequences. Taxon 55, 95–106. doi:10.2307/25065531

Wagstaff, S.J., Dawson, M.I., Venter, S., Munzinger, J., Crayn, D.M., Steane, D.A., Lemson, K.L., 2010. Origin, diversification, and classification of the Australasian genus Dracophyllum (Richeeae, Ericaceae). Ann. Missouri Bot. Gard. 97, 235–258. doi:10.3417/2008130

Weitemier, K., Straub, S. C., Cronn, R. C., Fishbein, M., Schmickl, R., McDonnell, A., & Liston, A. (2014). Hyb-Seq: Combining target enrichment and genome skimming for plant phylogenomics. Applications in plant sciences, 2(9), 1400042.

WCSP (6^th^ July, 2022). World Checklist of Selected Plant Families. Facilitated by the Royal Botanic Gardens, Kew. http://wcsp.science.kew.org/ Retrieved 2022, 6^th^ July.

Weising, K., Nybom, H., Pfenninger, M., Wolff, K., Kahl, G., 2005. DNA fingerprinting in plants: principles, methods, and applications 2nd. ed. CRC Press, BocaRaton, Florida.

Yu, Y., Harris, A.J., Blair, C., He, X., 2015. RASP (Reconstruct Ancestral State in Phylogenies): A tool for historical biogeography. Mol. Phylogenet. Evol. 87, 46–49. doi:10.1016/J.YMPEV.2015.03.008

Yule, G.U., 1925. A Mathematical Theory of Evolution, Based on the Conclusions of Dr. J. C. Willis, F.R.S. Philos. Trans. R. Soc. B Biol. Sci. 213, 21–87. doi:10.1098/rstb.1925.0002

Zahirovic, S., Matthews, K. J., Flament, N., Müller, R. D., Hill, K. C., Seton, M., & Gurnis, M. (2016). Tectonic evolution and deep mantle structure of the eastern Tethys since the latest Jurassic. Earth-Science Reviews, 162, 293–337.

Zuckerkandl, E., Pauling, L., 1965. Molecules as documents of evolutionary history. J. Theor. Biol. 8, 357–366. doi:10.1016/0022-5193(65)90083-4

