## Supplementary material for "Plastid phylogenomics clarifies broad-level relationships in *Bulbophyllum* (Orchidaceae) and provides insights into range evolution of Australasian section *Adelopetalum*"

**Supplementary material S1: Plant material used in this study with voucher details. Accessions included in divergence age estimations are indicated with an asterisk.**

| Taxon | Section | Voucher details | DNA number | Markers |
|---|---|---|---|---|
| *Bulbophyllum sigaldiae* Guillaumin (1955) | *Acrochaene* (Lindl.) J.J.Verm. Schuit. & de Vogel (2014) | CANB Orchid Research Group 7679 | CNS_G07581* | 70 plastid genes\|ITS_ETS cistron |
| *Bulbophyllum argyropus* 1 (Endl.) Rchb.f. (1876) | *Adelopetalum* (Fitzg.) J.J.Verm. (1993) | CANB Orchid Research Group 7052 | CNS_G03566* | 70 plastid genes\|ITS_ETS cistron |
| *Bulbophyllum argyropus* 2 (Endl.) Rchb.f. (1876) | *Adelopetalum* (Fitzg.) J.J.Verm. (1993) | CANB Sinclair, D. 5613 | CNS_G00333 | ITS\|matK\|ycf1 |
| *Bulbophyllum boonjee* 1 B.Gray & D.L.Jones (1984) | *Adelopetalum* (Fitzg.) J.J.Verm. (1993) | CANB Gray, B. 9761 | CNS_G07175* | 70 plastid genes\|ITS_ETS cistron |
| *Bulbophyllum boonjee* 2 B.Gray & D.L.Jones (1984) | *Adelopetalum* (Fitzg.) J.J.Verm. (1993) | CANB Jones, D.L. 4226 | CNS_G03563 | ITS\|matK\|ycf1 |
| *Bulbophyllum bracteatum* 1 (Fitzg.) F.M.Bailey (1891) | *Adelopetalum* (Fitzg.) J.J.Verm. (1993) | CNS Simpson, L. 497 | CNS_G01553* | 70 plastid genes\|ITS_ETS cistron |
| *Bulbophyllum bracteatum* 2 (Fitzg.) F.M.Bailey (1891) | *Adelopetalum* (Fitzg.) J.J.Verm. (1993) | CANB Orchid Research Group 5159 | CNS_G00383 | ITS\|matK\|ycf1 |
| **Taxon** | **Section** | **Voucher details** | **DNA number** | **Markers** |

| | | | | |
|---|---|---|---|---|
| *Bulbophyllum corythium* N.Hall (1981) | *Adelopetalum* (Fitzg.) J.J.Verm. (1993) | CANB Clements, M.A. 11238 | CNS_G00370* | ITS ycf1 |
| *Bulbophyllum elisae* 1 (F.Muell.) Benth. (1871) | *Adelopetalum* (Fitzg.) J.J.Verm. (1993) | CNS Simpson, L. 499 | CNS_G01549 | ITS\|matK\|ycf1 |
| *Bulbophyllum elisae* 2 (F.Muell.) Benth. (1871) | *Adelopetalum* (Fitzg.) J.J.Verm. (1993) | CNS Simpson, L. 500 | CNS_G01550* | ITS\|matK\|ycf1 |
| *Bulbophyllum elisae* 3 (F.Muell.) Benth. (1871) | *Adelopetalum* (Fitzg.) J.J.Verm. (1993) | CNS Simpson, L. 498 | CNS_G01548 | ITS\|matK\|ycf1 |
| *Bulbophyllum exiguum* 1 F.Muell. (1860) | *Adelopetalum* (Fitzg.) J.J.Verm. (1993) | CANB 667983 | CNS_G00322 | ITS\|matK\|ycf1 |
| *Bulbophyllum exiguum* 2 F.Muell. (1860) | *Adelopetalum* (Fitzg.) J.J.Verm. (1993) | CNS Simpson, L. 494 | CNS_G01558* | ITS\|matK\|ycf1 |
| *Bulbophyllum exiguum* 3 F.Muell. (1860) | *Adelopetalum* (Fitzg.) J.J.Verm. (1993) | CANB Schulte, K. 121 | CNS_G00234 | ITS\|matK\|ycf1 |
| *Bulbophyllum lageniforme* 1 F.M.Bailey (1904) | *Adelopetalum* (Fitzg.) J.J.Verm. (1993) | CNS Simpson, L. LS167B | CNS_G03707* | 70 plastid genes\|ITS_ETS cistron |
| *Bulbophyllum lageniforme* 2 F.M.Bailey (1904) | *Adelopetalum* (Fitzg.) J.J.Verm. (1993) | CNS Simpson, L. 180A | CNS_G04503 | ITS\|matK\|ycf1 |
| *Bulbophyllum lageniforme* 3 F.M.Bailey (1904) | *Adelopetalum* (Fitzg.) J.J.Verm. (1993) | CNS Schulte, K. 95 | CNS_G01630 | ITS\|matK\|ycf1 |
| *Bulbophyllum lilianiae* 1 Rendle (1917) | *Adelopetalum* (Fitzg.) J.J.Verm. (1993) | CNS Simpson, L. 337E | CNS_G07754* | 70 plastid genes\|ITS_ETS cistron |
| *Bulbophyllum lilianiae* 2 Rendle (1917) | *Adelopetalum* (Fitzg.) J.J.Verm. (1993) | CNS Simpson, L. 496 | CNS_G01561 | ITS\|matK\|ycf1 |
| *Bulbophyllum lilianiae* 3 Rendle (1917) | *Adelopetalum* (Fitzg.) J.J.Verm. (1993) | CNS Schulte, K. 87 | CNS_G01632 | ITS\|matK\|ycf1 |
| *Bulbophyllum lingulatum* Rendle | *Adelopetalum* (Fitzg.) J.J.Verm. (1993) | CANB Clements, M.A. 7959 | CNS_G06046* | 70 plastid genes\|ITS_ETS cistron |
| *Bulbophyllum newportii* 1 (F.M.Bailey) Rolfe (1909) | *Adelopetalum* (Fitzg.) J.J.Verm. (1993) | CNS Simpson, L. 155E | CNS_G03754* | 70 plastid genes\|ITS_ETS cistron |
| *Bulbophyllum newportii* 2 (F.M.Bailey) Rolfe (1909) | *Adelopetalum* (Fitzg.) J.J.Verm. (1993) | CNS Simpson, L. 495 | CNS_G01564 | ITS\|matK\|ycf1 |
| *Bulbophyllum newportii* 3 (F.M.Bailey) Rolfe (1909) | *Adelopetalum* (Fitzg.) J.J.Verm. (1993) | CNS Simpson, L. 492 | CNS_G01562 | ITS\|matK\|ycf1 |
| **Taxon** | **Section** | **Voucher details** | **DNA number** | **Markers** |

| | | | | |
|---|---|---|---|---|
| *Bulbophyllum tuberculatum* 1 Colenso | *Adelopetalum* (Fitzg.) J.J.Verm. (1993) | CANB Molloy, B.P.J. 112/99 | CNS_G03993 | ITS\|matK\|ycf1 |
| *Bulbophyllum tuberculatum* 2 Colenso | *Adelopetalum* (Fitzg.) J.J.Verm. (1993) | CHR 572204 | CHR572204* | ITS\|matK\|ycf1 |
| *Bulbophyllum weinthalii* ssp. *weinthalii* 1 R.S.Rogers (1933) | *Adelopetalum* (Fitzg.) J.J.Verm. (1993) | BRI M.T. Mathieson 730 | CNS_G06024* | 70 plastid genes\|ITS_ETS cistron |
| *Bulbophyllum weinthalii* ssp. *weinthalii* 2 R.S.Rogers (1933) | *Adelopetalum* (Fitzg.) J.J.Verm. (1993) | CNS Simpson, L. 493 | CNS_G01624 | ITS\|matK\|ycf1 |
| *Bulbophyllum weinthalii* ssp. *striatum* R.S. Rogers | *Adelopetalum* (Fitzg.) J.J.Verm. (1993) | CANB Orchid Research Group 7186 | CNS_G03564 | ITS\|matK\|ycf1 |
| *Bulbophyllum occlusum* Ridl., J. (1885) | *Alcistachys* Schltr. (1924) | CANB Orchid Reseach Group 3376 | CNS_G07291* | 70 plastid genes\|ITS_ETS cistron |
| *Bulbophyllum gymnopus* Hook.f. | *Altisceptrum* J.J.Sm. 1914 | CANB Orchid Research Group 7179 | CNS_G05207* | 70 plastid genes\|ITS_ETS cistron |
| *Bulbophyllum beccarii* Rchb.f. (1879) | *Beccariana* Pfitz. (1889) | CANB Orchid Research Group 7678 | CNS_G07559* | 70 plastid genes\|ITS_ETS cistron |
| *Bulbophyllum cruentum* Garay, Hamer & Siegerist (1992) | *Beccariana* Pfitz. (1889) | CANB Orchid Research Group 5524 | CNS_G06039* | 70 plastid genes\|ITS_ETS cistron |
| *Bulbophyllum elevatopunctatum* J.J.Sm. (1920) | *Beccariana* Pfitz. (1889) | CANB Orchid Research Group 7047 | CNS_G07187 | ITS_ETS cistron |
| *Bulbophyllum ericssonii* Kraenzl. (1893) | *Beccariana* Pfitz. (1889) | CANB Orchid Research Group 6981 | CNS_G05214* | 70 plastid genes\|ITS_ETS cistron |
| *Bulbophyllum foetidum* Schltr. | *Beccariana* Pfitz. (1889) | CANB Clements, M.A. 6431 | CNS_G05873* | 70 plastid genes\|ITS_ETS cistron |
| *Bulbophyllum uniflorum* (Blume) Hassk. (1844) | *Beccariana* Pfitz. (1889) | CANB Orchid Research Group 7670 | CNS_G07571* | 70 plastid genes\|ITS_ETS cistron |
| *Bulbophyllum wakoi* Howcroft (1999) | *Beccariana* Pfitz. (1889) | CANB Orchid Research Group 7176 | CNS_G05222 | ITS_ETS cistron |
| *Bulbophyllum biflorum* Teijsm. & Binn. (1854) | *Biflorae* Garay, Hamer & Siegrist (1994) | CANB Orchid Research Group 5887 | CNS_G05442* | 70 plastid genes\|ITS_ETS cistron |
| *Bulbophyllum lasiochilum* C.S.P.Parish & Rchb.f. (1874) | *Brachyantha* Rchb.f 1861 | CANB Clements, M.A. 7343 | CNS_G07204* | 70 plastid genes\|ITS_ETS cistron |
| **Taxon** | **Section** | **Voucher details** | **DNA number** | **Markers** |

| | | | | |
|---|---|---|---|---|
| *Bulbophyllum guttulatum* (Hook.f.) N.P.Balakr. (1970) | *Brachyantha* Rchb.f. (1861) | CANB Orchid Research Group 5587 | CNS_G05250* | 70 plastid genes\|ITS_ETS cistron |
| *Bulbophyllum macraei* (Lindl.) Rchb.f. | *Brachyantha* Rchb.f. (1861) | CANB Clements, M.A. 12421 | CNS_G06045* | 70 plastid genes\|ITS_ETS cistron |
| *Bulbophyllum nematopodum* F.Muell. (1872) | *Brachypus* Schlechter 1913 | CNS Harrison, Michael | CNS_G01612* | ITS\|ycf1 |
| *Bulbophyllum lineolatum* Schltr. (1913) | *Brachypus* Schltr. (1913) | CANB M.A. Clements 9580 | CNS_G05262* | 70 plastid genes\|ITS_ETS cistron |
| *Bulbophyllum maxillarioides* Schltr. (1905) | *Brachypus* Schltr. (1913) | CANB Clements, M.A. 7253 | CNS_G07332* | 70 plastid genes\|ITS_ETS cistron |
| *Bulbophyllum evasum* T.E.Hunt & Rupp (1950) | *Brachystachyae* Benth. & Hook.f. (1883). | CANB 599819 | CNS_G01247* | 70 plastid genes\|ITS_ETS cistron |
| *Bulbophyllum longissimum* (Ridl.) J.J.Sm. (1912) | *Cirrhopetaloides* Garay, Hamer & Siegerist (1994) | CANB Orchid Research Group 5525 | CNS_G06041* | 70 plastid genes\|ITS_ETS cistron |
| *Bulbophyllum putidum* (Teijsm. & Binn.) J.J.Sm. | *Cirrhopetaloides* Garay, Hamer & Siegerist (1994) | CANB Wallace, B. 19/91 | CNS_G05872* | 70 plastid genes\|ITS_ETS cistron |
| *Bulbophyllum forrestii* Seidenf. (1974) | *Cirrhopetalum* (Lindl.) Rchb.f. (1861) | CANB Orchid Research Group 1271 | CNS_G06037* | 70 plastid genes\|ITS_ETS cistron |
| *Bulbophyllum longiflorum* Thouars (1822) | *Cirrhopetalum* (Lindl.) Rchb.f. (1861) | CANB Orchid Research Group 7758 | CNS_G06022* | 70 plastid genes\|ITS_ETS cistron |
| *Bulbophyllum alkmaarense* J.J.Sm. (1911) | *Codonosiphon* Schlechter 1913 | CANB G. McCraith 076c,s4192 | CNS_G05454* | 70 plastid genes\|ITS_ETS cistron |
| *Bulbophyllum cruciatum* J.J.Sm. (1911) | *Codonosiphon* Schltr. (1911) | CBG 8600516 | CNS_G01056* | 70 plastid genes\|ITS_ETS cistron |
| *Bulbophyllum cauliflorum* Hook.f. (1890) | *Desmosanthes* (Blume) J.J.Sm. (1933) | CANB B. Wallace BJW 20/91 | CNS_G05243* | 70 plastid genes\|ITS_ETS cistron |
| *Bulbophyllum medusae* (Lindl.) Rchb.f. (1861) | *Desmosanthes* (Blume) J.J.Sm. (1933) | CANB Orchid Research Group 7684 | CNS_G07573* | 70 plastid genes\|ITS_ETS cistron |
| *Bulbophyllum pleurothallidanthum* Garay (1999) | *Desmosanthes* (Blume) J.J.Sm. (1933) | CANB Orchid Research Group 7681 | CNS_G07567* | 70 plastid genes\|ITS_ETS cistron |
| *Bulbophyllum gracillimum* (Rolfe) Rolfe (1907) | *Ephippium* Schlechter 1913 | CNS Field, A. | CNS_G01627* | ITS\|matK\|ycf1 |
| **Taxon** | **Section** | **Voucher details** | **DNA number** | **Markers** |

| | | | | |
|---|---|---|---|---|
| *Bulbophyllum haniffii* Carr (1932) | *Epicranthes* (Blume) Benth. & Hook.f. (1883) | CANB Orchid Research Group 7677 | CNS_G07566* | 70 plastid genes\|ITS_ETS cistron |
| *Bulbophyllum lindleyanum* Griff. (1851) | *Hirtula* Ridl. (1908), fide J.J.Verm. (2002) | CANB Orchid Research Group 6237 | CNS_G06035* | 70 plastid genes\|ITS_ETS cistron |
| *Bulbophyllum baladeanum* J.J.Sm. (1912) | *Hoplandra* J.J.Verm. (2008) | CANB M.A. Clements 11187 | CNS_G05443* | 70 plastid genes\|ITS_ETS cistron |
| *Bulbophyllum antenniferum* (Lindl.) Rchb.f. (1861) | *Hyalosema* (Rolfe) Schltr. (1911) | CANB T.Reeve 638 | CNS_G05294* | 70 plastid genes\|ITS_ETS cistron |
| *Bulbophyllum infundibuliforme* J.J.Sm. (1906) | *Hymenobractea* Schltr. (1913) | CANB Clements, M.A. 6616 | CNS_G07305 | ITS_ETS cistron |
| *Bulbophyllum digoelense* J.J.Sm. (1911) | *Intervallatae* Ridl. (1897) | CANB Orchid Research Group 6246 | CNS_G05276 | ITS_ETS cistron |
| *Bulbophyllum roseopictum* J.J.Verm., Schuit. & de Vogel | *Ione* [Lindley] J J Verm Schuit & de Vogel 2014 | CANB Clements, M.A 12085 | CNS_G03907* | 70 plastid genes\|ITS_ETS cistron |
| *Bulbophyllum lemniscatoides* Rolfe (1890) | *Lemniscata* Pfitz. (1888) | CANB Orchid Research Group 7153 | CNS_G05204* | 70 plastid genes\|ITS_ETS cistron |
| *Bulbophyllum levyae* Garay, Hamer & Siegerist (1995) | *Lepidorhiza* Schlechter 1911 | CANB Orchid Research Group 7674 | CNS_G07565* | 70 plastid genes\|ITS_ETS cistron |
| *Bulbophyllum echinolabium* J.J.Sm. (1934) | *Lepidorhiza* Schltr. (1911) | CANB G. McCraith 165 | CNS_G05237 | ITS_ETS cistron |
| *Bulbophyllum ovalifolium* (Blume) Lindl. (1830) | *Macrocaulia* (Blume) Aver. (1994) | CANB Orchid Research Group 6254 | CNS_G05210* | 70 plastid genes\|ITS_ETS cistron |
| *Bulbophyllum kaniense* Schltr. (1913) | *Macrouris* Schltr. (1913) | CANB M.A. Clements 6249 | CNS_G05254* | 70 plastid genes\|ITS_ETS cistron |
| *Bulbophyllum maximum* (Lindl.) Rchb.f. (1861) | *Megaclinium* (Lindl.) Summerh (1921) | CANB Orchid Research Group 6250 | CNS_G05289* | 70 plastid genes\|ITS_ETS cistron |
| *Bulbophyllum globuliforme* 1 Nicholls (1938) | *Minutissima* Pftiz. (1888) | CNS Simpson, L. 501 | CNS_G01584 | ITS\|matK\|ycf1 |
| *Bulbophyllum globuliforme* 2 Nicholls (1938) | *Minutissima* Pftiz. (1888) | CANB McCraith, G. 86 | CNS_G00392* | ITS\|matK\|ycf1 |
| *Bulbophyllum keekee* N.Hall (1977) | *Minutissima* Pftiz. (1888) | CANB Clements, M.A. 5675 | CNS_G07329* | 70 plastid genes\|ITS_ETS cistron |
| **Taxon** | **Section** | **Voucher details** | **DNA number** | **Markers** |

| | | | | |
|---|---|---|---|---|
| *Bulbophyllum minutissimum* 1 (F.Muell.) F.Muell. (1878) | *Minutissima* Pftiz. (1888) | CNS Simpson, L. 503 | CNS_G01587* | 70 plastid genes\|ITS_ETS cistron |
| *Bulbophyllum minutissimum* 2 (F.Muell.) F.Muell. (1878) | *Minutissima* Pftiz. (1888) | CNS Simpson, L. 502 | CNS_G01585 | ITS\|matK\|ycf1 |
| *Bulbophyllum moniliforme* E.C.Parish & Rchb.f. | *Minutissima* Pftiz. (1888) | CANB Clements, M.A. 9605 | CNS_G03763* | 70 plastid genes\|ITS_ETS cistron |
| *Bulbophyllum mucronatum* (Blume) Lindl. (1830) | *Minutissima* Pftiz. (1888) | CANB M.A. Clements MAC 12379 | CNS_G05440* | 70 plastid genes\|ITS_ETS cistron |
| *Bulbophyllum pygmaeum* 1 (Sm.) Lindl. | *Minutissima* Pftiz. (1888) | CANB Molloy, B.P.J. 062/98 | CNS_G04599* | 70 plastid genes\|ITS_ETS cistron |
| *Bulbophyllum pygmaeum* 2 (Sm.) Lindl. | *Minutissima* Pftiz. (1888) | CANB Molloy, B.P.J. 134/99 | CNS_G03988* | ITS\|matK\|ycf1 |
| *Bulbophyllum ciliatum* (Blume) Lindl. (1830) | *Monanthaparva* Ridl. 1896 | CANB Orchid Research Group 4316 | CNS_G05450* | 70 plastid genes\|ITS_ETS cistron |
| *Bulbophyllum dischidiifolium* J.J.Sm. (1909) | *Monanthes* (Blume) Aver. (1994) | CBG 9013189 | CNS_G05278 | 70 plastid genes\|ITS_ETS cistron |
| *Bulbophyllum macphersonii* Rupp (1934) | *Monanthes* (Blume) Aver. (1994) | CANB Schulte, K. 85 CBG8600516 | CNS_G01039* | ITS\|ycf1 |
| *Bulbophyllum clandestinum* 1 Lindl. (1841) | *Oxysepala* Bentham & J D Hook.f. 1883 | CANB Reeve, T.M. 735 | CNS_G07307* | 70 plastid genes\|ITS_ETS cistron |
| *Bulbophyllum clandestinum* 2 Lindl. (1841) | *Oxysepala* Bentham & J D Hook.f. 1883 | CANB Orchid Research Group 3080 | CNS_G05272* | 70 plastid genes\|ITS_ETS cistron |
| *Bulbophyllum gadgarrense* Rupp (1949) | *Oxysepala* Bentham & J D Hook.f. 1883 | CANB Clements, M.A. 8535 | CNS_G00933* | ITS\|ycf1 |
| *Bulbophyllum grandimesense* B.Gray & D.L.Jones (1989) | *Oxysepala* Bentham & J D Hook.f. 1883 | CANB Roberts, L.J. s.n. | CNS_G01588* | ITS\|ycf1 |
| *Bulbophyllum lamingtonense* D.L.Jones (1993) | *Oxysepala* Bentham & J D Hook.f. 1883 | CANB Orchid Research Group 6946 | CNS_G01013* | ITS\|ycf1 |
| *Bulbophyllum lewisense* B.Gray & D.L.Jones (1989) | *Oxysepala* Bentham & J D Hook.f. 1883 | CNS Harrison, Michael | CNS_G01608* | ITS\|ycf1 |
| *Bulbophyllum rhopalophorum* Schltr. (1913) | *Oxysepala* Bentham & J D Hook.f. 1883 | CBG 9013094 | CNS_G05261* | 70 plastid genes\|ITS_ETS cistron |
| *Bulbophyllum schillerianum* Rchb.f. (1860) | *Oxysepala* Bentham & J D Hook.f. 1883 | CANB Forster, P. 24990 | CNS_G01597* | ITS\|ycf1 |
| **Taxon** | **Section** | **Voucher details** | **DNA number** | **Markers** |

| | | | | |
|---|---|---|---|---|
| *Bulbophyllum shepherdii* (F.Muell.) Rchb.f. (1871) | *Oxysepala* Bentham & J D Hook.f. 1883 | CANB Orchid Research Group 4858 | CNS_G00804* | ITS\|ycf1 |
| *Bulbophyllum wadsworthii* Dockrill (1964) | *Oxysepala* Bentham & J D Hook.f. 1883 | CNS Kilgour, C.D. 1143 | CNS_G00574* | ITS\|ycf1 |
| *Bulbophyllum windsorense* B.Gray & D.L.Jones (1989) | *Oxysepala* Bentham & J D Hook.f. 1883 | CNS Harrison, Michael | CNS_G01610* | ITS\|ycf1 |
| *Bulbophyllum pachypus* Schltr. (1924) | *Pachychlamys* Schltr. (1925) | CANB Clements, M.A. 12774 | CNS_G07740* | 70 plastid genes\|ITS_ETS cistron |
| *Bulbophyllum sauguetiense* Schltr. (1913) | *Papulipetalum* Schltr. (1913) | CANB M.A. Clements 7235 | CNS_G05258* | 70 plastid genes\|ITS_ETS cistron |
| *Bulbophyllum oreomene* J.J.Verm., Schuit. & de Vogel (2014) | *Pedilochilus* (Schltr.) J.J.Ver. & P.O'Byrne (2011) | Australian National University Hope, G. 10850 | CNS_G07517* | 70 plastid genes\|ITS_ETS cistron |
| *Bulbophyllum absconditum* J.J.Sm. (1905) | *Pelma* (Finet) Schltr. (1913) | CANB M.A. Clements MAC11242 | CNS_G05247* | 70 plastid genes\|ITS_ETS cistron |
| *Bulbophyllum triaristella* Schltr. (1913) | *Peltopus* Schlechter 1913 | CANB Clements, M.A. | CNS_G01202* | 70 plastid genes\|ITS_ETS cistron |
| *Bulbophyllum aphanopetalum* Schltr. (1906) | *Peltopus* Schltr. (1913) | CANB P. Ziesing | CNS_G05241* | 70 plastid genes\|ITS_ETS cistron |
| *Bulbophyllum baronii* Ridl. (1885) | *Ploiarium* Schlechter (1925) | CANB Orchid Research Group 6242 | CNS_G06040* | 70 plastid genes\|ITS_ETS cistron |
| *Bulbophyllum mirum* J.J.Sm. (1906) | *Plumata* J.J.Verm., Schuit. & de Vogel (2014) | CANB Orchid Research Group 5501 | CNS_G06031* | 70 plastid genes\|ITS_ETS cistron |
| *Bulbophyllum plumatum* Ames (1915) | *Plumata* J.J.Verm., Schuit. & de Vogel (2014) | CANB Orchid Research Group 7673 | CNS_G07578* | 70 plastid genes\|ITS_ETS cistron |
| *Bulbophyllum acutilingue* J.J.Sm.(1908) | *Polymeres* (Blume) J.J.Verm. & O'Byrne (2008) | CANB Rose, S. 25 | CNS_G07612* | 70 plastid genes\|ITS_ETS cistron |
| *Bulbophyllum bowkettiae* F.M.Bailey (1884) | *Polymeres* (Blume) J.J.Verm. & O'Byrne (2008) | CANB 596002 | CNS_G00826* | ITS\|ycf1 |
| *Bulbophyllum elassoglossum* Siegerist, Amer. (1991) | *Polymeres* (Blume) J.J.Verm. & O'Byrne (2008) | CANB M.A. Clements 11502 | CNS_G05456* | 70 plastid genes\|ITS_ETS cistron |
| **Taxon** | **Section** | **Voucher details** | **DNA number** | **Markers** |

| | | | | |
|---|---|---|---|---|
| *Bulbophyllum fruticicola* Schltr. (1905) | *Polymeres* (Blume) J.J.Verm. & O'Byrne (2008) | CANB Reeve, T.M. 3372 | CNS_G07481* | 70 plastid genes\|ITS_ETS cistron |
| *Bulbophyllum johnsonii* T.E.Hunt (1950) | *Polymeres* (Blume) J.J.Verm. & O'Byrne (2008) | CNS Kilgour, C.D. 150 | CNS_G00538* | ITS\|ycf1 |
| *Bulbophyllum maxillare* 1 (Lindl.) Rchb.f. (1861) | *Polymeres* (Blume) J.J.Verm. & O'Byrne (2008) | CANB D.L. Jones 3587 | CNS_G05267* | 70 plastid genes\|ITS_ETS cistron |
| *Bulbophyllum maxillare* 2 (Lindl.) Rchb.f. (1861) | *Polymeres* (Blume) J.J.Verm. & O'Byrne (2008) | CANB Taylor, J.M. 284 | CNS_G01299* | 70 plastid genes\|ITS_ETS cistron |
| *Bulbophyllum ocellatum* Cootes & M.A.Clem. (2011) | *Polymeres* (Blume) J.J.Verm. & O'Byrne (2008) | CANB M.A. Clements 11603 | CNS_G05293* | 70 plastid genes\|ITS_ETS cistron |
| *Bulbophyllum rhodoglossum Schltr.* (1913) | *Polymeres* (Blume) J.J.Verm. & O'Byrne (2008) | CANB Reeve, T.M. 995 | CNS_G07310* | 70 plastid genes\|ITS_ETS cistron |
| *Bulbophyllum wolfei* B.Gray & D.L.Jones (1991) | *Polymeres* (Blume) J.J.Verm. & O'Byrne (2008) | CANB Jones, D.L. 4353 | CNS_G01623* | ITS\|ycf1 |
| *Bulbophyllum radicans* F.M.Bailey (1897) | *Polymeres* Verm & O'Byrne 2008 | CANB R. lockyer 6 | CNS_G05228* | 70 plastid genes\|ITS_ETS cistron |
| *Bulbophyllum schinzianum* Kraenzl. (1899) | *Ptiloglossum* Lindl. (1862) | CANB Clements, M.A. 11554 | CNS_G01642* | 70 plastid genes\|ITS_ETS cistron |
| *Bulbophyllum intricatum* Seidenf. (1979) | *Racemosae* Benth. & Hook.f. (1883) | CANB G. McCraith 083 | CNS_G05202* | 70 plastid genes\|ITS_ETS cistron |
| *Bulbophyllum propinquum* Kraenzl. (1908) | *Racemosae* Benth. & Hook.f. (1883) | CANB Orchid Research Group 7667 | CNS_G07576* | 70 plastid genes\|ITS_ETS cistron |
| *Bulbophyllum saurocephalum* | *Saurocephalum* Schltr. (1912) | CANB M.A. Clements 11496 | CNS_G05212* | 70 plastid genes\|ITS_ETS cistron |
| *Bulbophyllum baileyi* F.Muell. (1875) | *Sestochilos* (Breda) Benth & Hook.f. (1883) | CANB Jones, D.L. 18707c | CNS_G00242* | ITS\|matK\|ycf1 |
| **Taxon** | **Section** | **Voucher details** | **DNA number** | **Markers** |

| | | | | |
|---|---|---|---|---|
| *Bulbophyllum gjellerupii* J.J.Sm. (1929) | *Sestochilos* (Breda) Benth & Hook.f. (1883) | CANB M.A. Clements 11594 | CNS_G05270* | 70 plastid genes\|ITS_ETS cistron |
| *Bulbophyllum lobbii* Lindl. (1847) | *Sestochilos* (Breda) Benth & Hook.f. (1883) | CANB Orchid Research Group 5166 | CNS_G05229* | 70 plastid genes\|ITS_ETS cistron |
| *Bulbophyllum macranthum* Lindl. (1844) | *Sestochilos* (Breda) Benth & Hook.f. (1883) | CANB 952179.1 | CNS_G05452 | ITS_ETS cistron |
| *Bulbophyllum flavescens* (Blume) Lindl. (1830) | *Stachysanthes* (Blume) Averyanov (1994) | CANB M.A. Clements 11548 | CNS_G05280* | 70 plastid genes\|ITS_ETS cistron |
| *Bulbophyllum cambodianum* (H.Wendl. & Kraenzl.) Rolfe (1897) | *Trias* (Lindl.) J.J.Verm., Schuit. & de Vogel (2014) | CANB R.Cowen 3077 | CNS_G05275* | 70 plastid genes\|ITS_ETS cistron |
| *Bulbophyllum tripudians* C.S.P.Parish & Rchb.f. (1875) | *Tripudianthes* Seidenf. (1979) | CANB Orchid Research Group 3081 | CNS_G05249* | 70 plastid genes\|ITS_ETS cistron |
| *Bulbophyllum cylindrobulbum* Schltr. (1905) | *Uncifera* Schltr. (1912) | CANB M.A. Clements MAC11200 | CNS_G05215* | 70 plastid genes\|ITS_ETS cistron |
| *Bulbophyllum sp.* | | CANB Reeve, T.M. 1097 | CNS_G07341 | ITS_ETS cistron |
| *Coelogyne flaccida* Lindl. | | WIS v0289172 | | plastid genes\|ITS |
| *Dendrobium brunneum* Schuit. & Peter B.Adams | | CANB Orchid Research Group 5485 | CNS_G00781* | plastid genes\|ITS |
| *Dendrobium cunninghamii* Lindl. | | CANB Molloy, B.P.J. 061/98 | CNS_G02336* | plastid genes\|ITS |
| *Dendrobium fuscescens* (Griff.) Summerh. | | CANB Orchid Research Group 6920 | CNS_G01021* | plastid genes\|ITS |
| *Dendrobium macropus* (Endl.) Rchb.f. | | CANB Ziesing, P.D. 345 | CNS_G02316* | plastid genes\|ITS |
| *Dendrobium moniliforme* (L.) Sw. | | CANB Clements, M.A. 8262 | CNS_G00882* | plastid genes\|ITS |
| *Dendrobium muricatum* Finet | | CANB Clements, M.A. 3093 | CNS_G02350* | plastid genes\|ITS |
| *Dienia ophrydis* (J.Koenig) Seidenf. | | CBG 9306410 | CNS_G01303* | plastid genes\|ITS |
| **Taxon** | **Section** | **Voucher details** | **DNA number** | **Markers** |

| | | | | |
|---|---|---|---|---|
| *Neottia Cordata* (L.) Rich. | | CANB Clements, M.A. 9865 | CNS_G05314* | plastid genes\|ITS |
| *Nervilia concolor* (Blume) Schltr. | | CANB Roberts, L. ORG3787 | CNS_G03442* | plastid genes\|ITS |
| *Oberonia complanata* (A.Cunn.) M.A.Clem. & D.L.Jones | | CNS135322 | CNS_G00309* | plastid genes\|ITS |

**Supplementary material S2: Plastid genes included in analyses.**

| | | | | | | |
|---|---|---|---|---|---|---|
| accD | infA | psaB | psbH | rpl2 | rpoB | rps14 |
| atpA | matK | psaC | psbI | rpl14 | rpoC1 | rps15 |
| atpB | orf42 | psaI | psbJ | rpl16 (in part) | rpoC2 | rps16 |
| atpE | petA | psaJ | psbK | rpl20 | rps2 | rps18 |
| atpF | petB (in part) | psbA | psbL | rpl22 | rps3 | rps19 |
| atpH | petD (in part) | psbB | psbM | rpl23 | rps4 | ycf1 |
| atpI | petG | psbC | psbN | rpl32 | rps7 | ycf2 |
| ccsA | petL | psbD | psbT | rpl33 | rps8 | ycf3 |
| cemA | petN | psbE | psbZ | rpl36 | rps11 | ycf4 |
| clpP | psaA | psbF | rbcL | rpoA | rps12 | ycf68 |

**Supplementary material S3: PCR reaction protocols.**

PCR reactions were carried out in 20 µL volumes. ITS reactions consisted of 2.5 µL PCR buffer (200mM Tris HCl pH 8.4, 500mM KCl), 0.5 µL MgCl (25mM), 0.5 µL each of forward and reverse primer (10µM), 0.5 µL dNTPs (10mM), 1 µL DMSO, 0.9 µL BSA, 0.4 µL KAPA Taq DNA polymerase (5U/µL) (Kapa Biosystems, Wilmington, USA), 10.7 µL $H_2O$ and 2.5 µL gDNA (ca. 5 ng/µL). For the amplification of *mat*K and *ycf*1, PCR reactions were carried out with 4 µL 5x High-Fidelity Buffer, 0.8 µL each of forward and reverse primer (10µM), 0.4µL dNTPs (10 mM), 0.6µL DMSO, 0.25 µL iProof High-Fidelity DNA polymerase (5U/µL), (Thermo Fisher Scientific, Waltham, USA), 11.15 µL $H_2O$, and 2 µL gDNA (ca. 5 ng/µL).

For ITS a touchdown PCR was carried out with an initial denaturation step at 94˚C for 2 min, followed by 7 cycles of 94 ˚C denaturation for 45 sec, 66 ˚C annealing for 45 sec (reducing 1 ºC at each cycle) and 72 ˚C extension for 1 min and 30 sec, with a final extension at 72 ˚C for 5 min. For *mat*K, an initial denaturation step was carried out at 98 ˚C for 45 sec followed by 35 cycles of 98 ˚C denaturation for 10 sec, 60 ˚C annealing for 45 sec and 72 ˚C extension/elongation for 45 sec, with a final extension at 72 ˚C for 10 min. For *ycf*1, PRC reactions were carried out with the same conditions as for *mat*K, except for the annealing temperature which was set to 63 ˚C. PCR products were cleaned using exonuclease (ExoI) and alkaline phosphatase (FastAP) (Thermo Fisher Scientific,

Waltham, USA) using 7.5µL of PCR product, 0.25µL of ExoI, 1 µL of FastAP and 1.25 µL of $H_2O$ incubated for 15 mins at 37 ˚C, followed by 15 mins at 85 ˚C.

**Supplementary material S4: Comparison of likelihood scores for divergence dating analyses with alternative clock and speciation and extinction models.**

| log file | AICM likelihood | +/- | burnin | bootstrap replicates | delta_AICM |
|---|---|---|---|---|---|
| relaxed log normal \| Birth Death | 477294.182 | 3.6993 | 0 | 1000 | |
| relaxed log normal \| Yule | 477309.447 | 3.9273 | 0 | 1000 | 15.2643 |
| strict \| Yule | 480530.16 | 2.6903 | 0 | 1000 | 3235.978 |
| strict \| Birth Death | 480531.156 | 2.4152 | 0 | 1000 | 3236.9736 |

**Supplementary material S5: Comparison of likelihood scores for three models of range evolution.**

| | LnL | numparams | d | e | AICc | AICc_wt |
|---|---|---|---|---|---|---|
| BAYAREALIKE | -287.2 | 2 | 0.0048 | 0.053 | 578.5 | 0.88 |
| DEC | -289.2 | 2 | 0.0079 | 0.0064 | 582.6 | 0.12 |
| DIVALIKE | -301.7 | 2 | 0.009 | 0.0033 | 607.6 | 4.40E-07 |

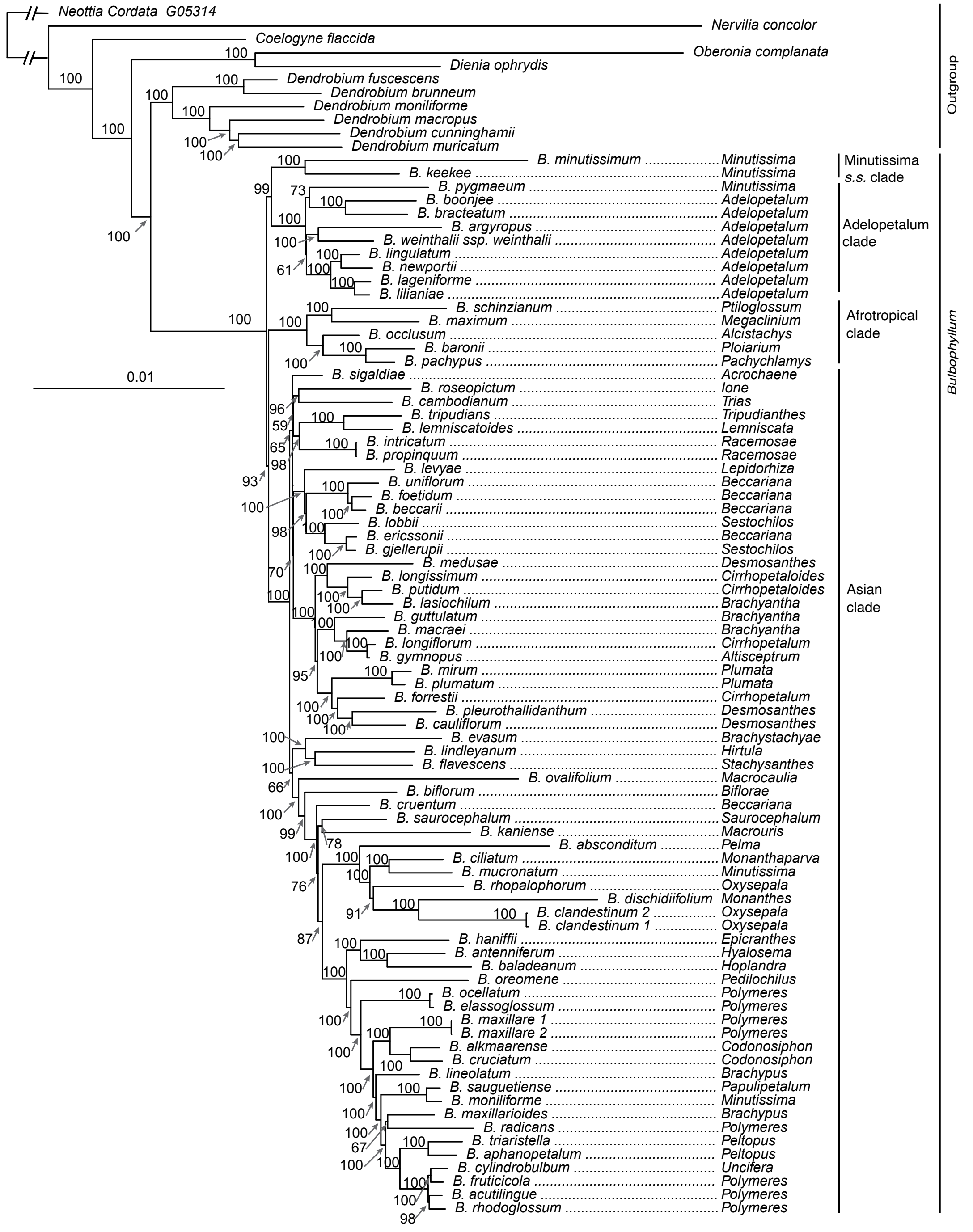


**Supplementary material S6: Maximum likelihood phylogenetic reconstruction of *Bulbophyllum* based on the 70 gene complete plastid dataset with reduced sampling.**

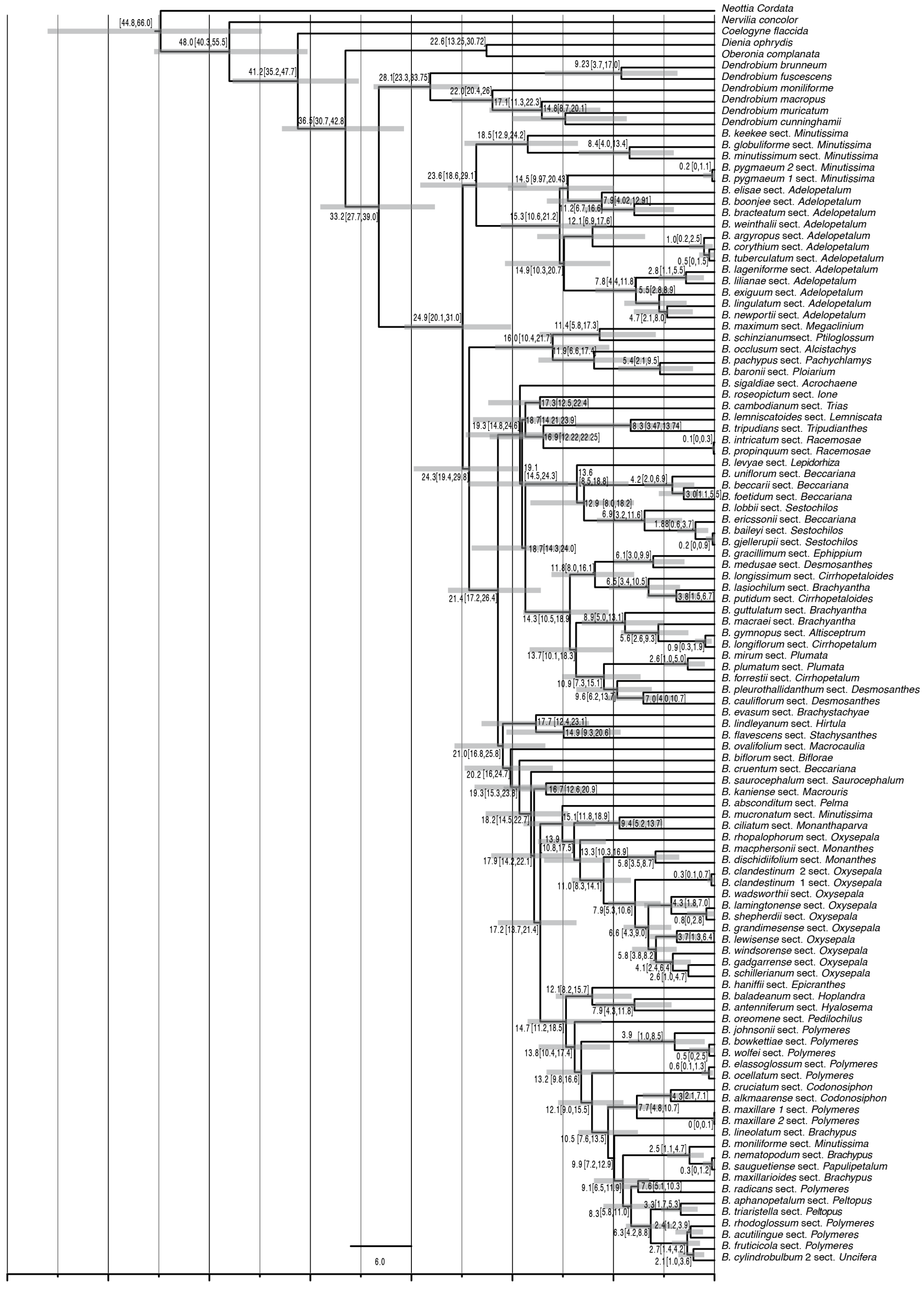

**Supplementary material S7: Maximum clade credibility chronogram for *Bulbophyllum* based on 70 plastid coding sequences, relaxed log normal clock and birth death prior. Divergence dates and 95% highest posterior density values are indicated adjacent to nodes. Grey bars indicate 95% highest posterior density.**

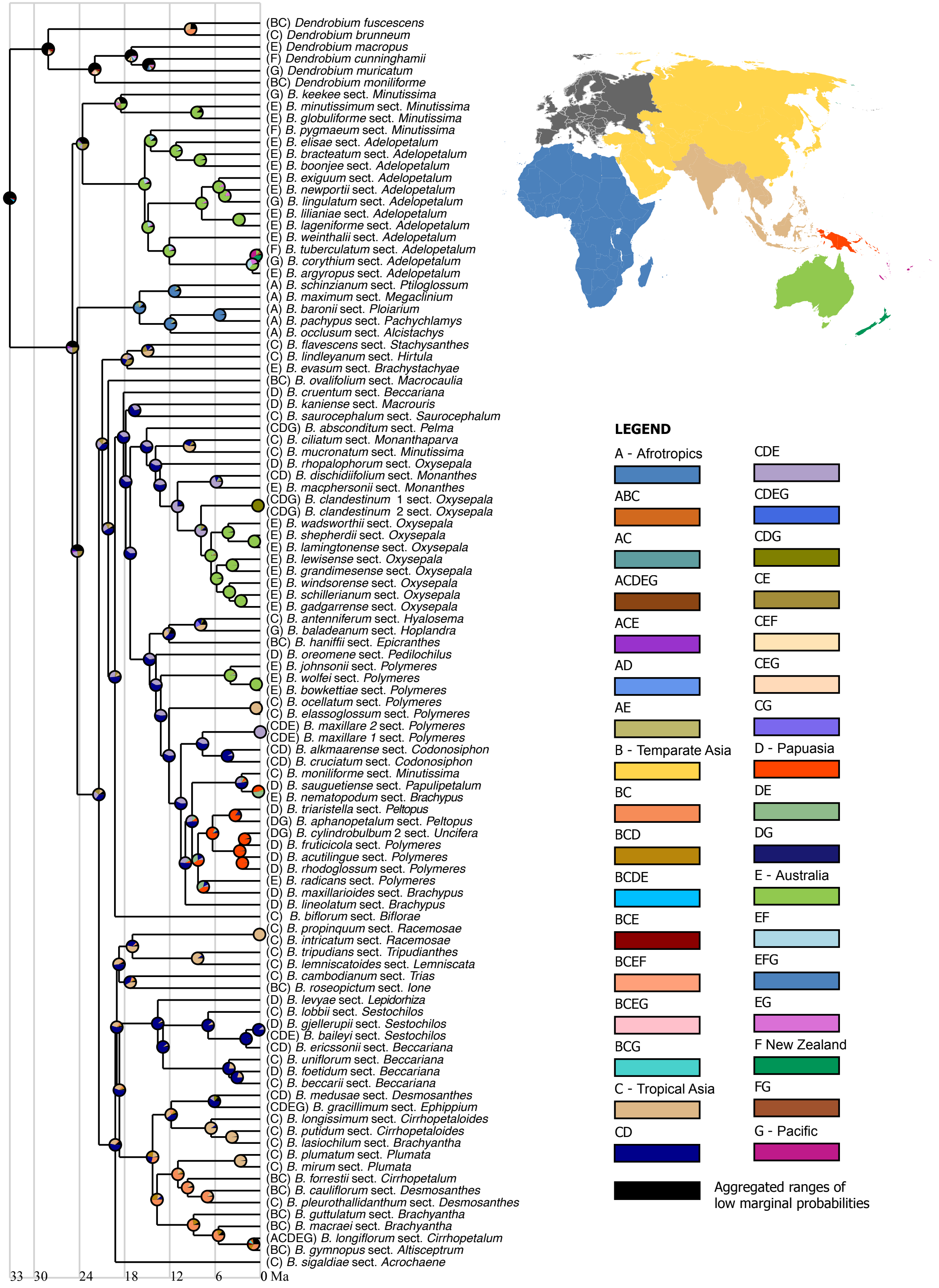


**Supplementary material S8: Ancestral area reconstruction based on a the BAYAREALIKE model with pie charts at internal nodes representing marginal probabilities for alternative ancestral areas. Map shows geographic regions delineated in the analysis and legend shows color-coded geographic regions and shared ancestral ranges.**

**Supplementary material S9: Range probabilities reconstructed for each node in the ancestral area reconstruction based on a the BAYAREALIKE model. Nodes IDs are presented in Supplementary Material S10.**

| Node ID | Range probabilities (%) |
|---|---|
| node 106 | BC 31; BCD 16; BCE 9; BCG 9; ABC 9; BCDE 4; BCDG 4; ABCD 4; BCEG 2; ABCE 2; ABCG 2; BCDEG 1; ABCDE 1; ABCDG 1; CD 1; ABCEG 1 |
| node 107 | BC 70; BCD 17; BCE 3; BCG 3; ABC 3; BCDE 1; BCDG 1; ABCD 1 |
| node 108 | BC 73; BCD 20; BCE 2; BCG 1; ABC 1; C 1 |
| node 109 | BC 85; C 12; BCD 2 |
| node 110 | BC 83; C 10; BCD 4; CD 1 |
| node 111 | C 97; BC 2 |
| node 112 | BC 74; C 15; BCD 7; CD 2 |
| node 113 | BC 50; BCD 28; C 11; CD 9; BCE 1 |
| node 114 | C 98; CD 1; BC 1 |
| node 115 | C 90; CD 6; BC 3 |
| node 116 | CD 60; CDE 14; CDG 13; BCD 4; CDEG 3; C 1; BCDE 1; BCDG 1 |
| node 117 | CD 40; C 31; BC 13; BCD 10; CE 1; CDE 1; CDG 1; CG 1; BCE 1 |
| node 118 | BC 28; CD 25; BCD 24; C 20; BCE 1; CE 1 |
| **Node ID** | **Range probabilities (%)** |

| | |
|---|---|
| node 119 | CD 80; C 19 |
| node 120 | CD 79; C 21 |
| node 121 | CD 89; CDE 10; D 1 |
| node 122 | CD 97; CDE 2 |
| node 123 | CD 89; C 8; CDE 1; BCD 1 |
| node 124 | CD 90; C 5; BCD 2; CDE 1 |
| node 125 | CD 89; C 5; BCD 3; CDE 1 |
| node 126 | CD 56; C 27; BC 7; BCD 6; CE 2; CDE 2 |
| node 127 | C 41; CD 39; BC 9; BCD 5; CE 2; CDE 1 |
| node 128 | C 90; CD 6; BC 2; CE 1 |
| node 129 | C 100 |
| node 130 | C 53; CD 35; BC 6; BCD 2; CE 2; CDE 1 |
| node 131 | CD 49; C 35; BC 7; BCD 5; CE 2; CDE 1 |
| node 132 | CD 55; C 28; BC 7; BCD 5; CE 2; CDE 2 |
| node 133 | CD 54; C 28; BC 6; BCD 5; CE 3; CDE 3 |
| node 134 | D 42; DE 37; CD 15; CDE 6 |
| **Node ID** | **Range probabilities (%)** |

| | |
|---|---|
| node 135 | D 99; DG 1 |
| node 136 | D 95; DG 5 |
| node 137 | D 98; DG 2 |
| node 138 | D 84; DG 13; DE 1; CD 1 |
| node 139 | D 83; DE 8; CD 5; DG 3 |
| node 140 | D 41; DE 33; CD 18; CDE 7 |
| node 141 | DE 49; D 45; E 6 |
| node 142 | CD 49; CDE 25; DE 9; D 9; CE 6; C 2; E 1 |
| node 143 | CD 47; CDE 28; DE 14; D 11 |
| node 144 | CD 51; CDE 32; DE 9; D 8 |
| node 145 | CD 92; CDE 7 |
| node 146 | CDE 100 |
| node 147 | CD 58; CDE 41 |
| node 148 | CD 58; CDE 41; DE 1 |
| node 149 | C 100 |
| node 150 | CD 57; CDE 41 |
| **Node ID** | **Range probabilities (%)** |

| | |
|---|---|
| node 151 | E 100 |
| node 152 | E 88; CE 5; DE 5 |
| node 153 | CD 54; CDE 44 |
| node 154 | CD 59; CDE 39 |
| node 155 | C 42; CG 22; CD 15; CDG 6; CE 5; CEG 2; BC 2; CDE 1; BCG 1; CDEG 1; BCD 1 |
| node 156 | CD 38; C 25; CDE 10; CE 8; CDG 4; BCD 3; CG 3; BC 2; CDEG 1; CEG 1; BCDE 1; BCE 1 |
| node 157 | CD 61; CDE 36; CDG 1; BCD 1 |
| node 158 | E 100 |
| node 159 | E 99 |
| node 160 | E 99 |
| node 161 | E 96; DE 2; CE 2 |
| node 162 | E 100 |
| node 163 | E 97; DE 2; CE 2 |
| node 164 | E 85; DE 7; CE 7; CDE 1 |
| node 165 | CDG 100 |
| node 166 | CDE 62; DE 13; CE 13; CD 5; CDEG 2; E 1; DEG 1; CEG 1 |
| **Node ID** | **Range probabilities (%)** |

| | |
|---|---|
| node 167 | CDE 70; CD 13; DE 7; CE 6 |
| node 168 | CDE 76; CD 16; DE 4; CE 3; CDEG 1 |
| node 169 | CD 54; CDE 43; DE 1; CDG 1; D 1 |
| node 170 | C 56; CD 28; CE 9; CDE 5 |
| node 171 | CD 58; CDE 40; CDG 1 |
| node 172 | CD 60; CDE 38; CDG 1; CDEG 1 |
| node 173 | CD 62; CDE 37 |
| node 174 | CD 67; CDE 29; BCD 1 |
| node 175 | CD 62; CDE 36; BCD 1 |
| node 176 | CD 62; CDE 36; BCD 1 |
| node 177 | CD 49; CDE 28; CE 10; C 8; BCD 2; BCDE 1; BC 1; BCE 1 |
| node 178 | CD 42; CDE 25; CE 15; C 10; BCD 3; BC 2; BCE 1; BCDE 1 |
| node 179 | C 45; CE 24; CD 18; CDE 6; BC 2; BCE 1; BCD 1; AC 1 |
| node 180 | CE 32; CD 23; C 22; CDE 15; BC 2; BCE 2; BCD 1; BCDE 1 |
| node 181 | CD 37; CDE 25; CE 19; C 11; BC 2; BCD 2; BCE 2; BCDE 1 |
| node 182 | CD 35; CDE 24; CE 19; C 11; BC 3; BCD 2; BCE 2; BCDE 1; ACD 1 |
| **Node ID** | **Range probabilities (%)** |

| | |
|---|---|
| node 183 | A 97; AC 1; AE 1; AD 1 |
| node 184 | A 81; AC 6; AE 5; AD 3; AG 1; AB 1; AF 1 |
| node 185 | A 80; AC 6; AE 6; AD 3; AG 1; AB 1; AF 1 |
| node 186 | A 57; AC 13; AE 11; AD 6; ACE 2; AG 2; AB 1; ACD 1; AF 1; ADE 1 |
| node 187 | CE 24; CDE 20; ACE 8; CD 7; ACDE 4; CEG 3; BCE 3; C 3; ACD 2; DE 2; AC 2; CDEG 2; CEF 2; BCDE 1; E 1; AE 1; CDEF 1; ADE 1; CDG 1; BC 1; ABCE 1; CG 1; ACEG 1; BCD 1; CDF 1 |
| node 188 | F 41; G 34; E 11; FG 9; EF 3; EG 2 |
| node 189 | E 36; EF 33; EG 26; EFG 3; F 2; G 1 |
| node 190 | E 72; EF 16; EG 7; CE 1; DE 1; EFG 1; AE 1 |
| node 191 | E 99 |
| node 192 | E 85; EG 15 |
| node 193 | E 90; EG 9 |
| node 194 | E 92; EG 6; EF 1 |
| node 195 | E 72; EF 16; EG 6; CE 2; DE 1; AE 1 |
| node 196 | E 95; EF 2; EG 1; CE 1 |
| node 197 | E 89; EF 6; EG 2; CE 1; DE 1 |
| node 198 | E 71; EF 18; EG 5; CE 2; DE 1; AE 1 |
| **Node ID** | **Range probabilities (%)** |

| | |
|---|---|
| node 199 | E 71; EF 16; EG 6; CE 3; DE 1; AE 1; EFG 1 |
| node 200 | E 82; CE 5; EG 4; DE 3; AE 2; EF 1; BE 1 |
| node 201 | E 24; CE 18; EG 13; DE 7; CEG 6; CDE 4; AE 4; DEG 3; ACE 2; EF 2; BE 2; CDEG 1; AEG 1; CEF 1; BCE 1; ADE 1; ACEG 1; EFG 1; BEG 1 |
| node 202 | CE 26; CDE 13; E 11; DE 7; ACE 6; CEG 5; AE 3; EG 3; CEF 3; BCE 2; ACDE 2; CDEG 2; ADE 1; DEG 1; CDEF 1; BE 1; EF 1; ACEG 1; BCDE 1; DEF 1; AEG 1 |
| node 203 | CE 26; CDE 21; ACE 8; CEG 5; ACDE 4; BCE 3; CDEG 3; DE 3; CEF 2; CD 2; CDEF 2; BCDE 1; E 1; AE 1; ADE 1; ACD 1; ACEG 1; AC 1; ABCE 1; C 1; CDG 1; DEG 1; ACEF 1; EG 1 |
| node 204 | EG 8; FG 8; EF 7; EFG 7; G 4; CEG 4; F 4; CEF 4; CFG 3; CE 3; E 3; CG 3; CF 3; CEFG 3; BEG 2; BEF 2; BFG 2; BG 2; BE 2; BF 2; BEFG 2; BCE 1; BCG 1; BCEG 1; C 1; BCF 1; BCEF 1; BCFG 1; BCEFG 1; DEG 1; DEF 1; BC 1; DFG 1; AEG 1; AEF 1; DG 1; DEFG 1; DF 1; DE 1; AFG 1 |
| node 205 | EG 8; EF 7; EFG 6; CEG 6; CE 5; CEF 5; E 4; CEFG 4; BEG 3; FG 3; BEF 3; BE 3; BCE 2; CFG 2; CG 2; CF 2; BCEG 2; BEFG 2; BCEF 2; G 1; BCEFG 1; F 1; BFG 1; BG 1; BCG 1; BF 1; BCF 1; DEG 1; DEF 1; BCFG 1; BC 1; DE 1; CDE 1; AEG 1; C 1; AEF 1; CDEG 1; CDEF 1; AE 1; ACE 1; DEFG 1 |
| node 206 | BCE 8; CE 7; CEG 6; CEF 6; BCEG 6; BCEF 5; CEFG 3; BCEFG 3; BCG 3; BCF 2; BC 2; CG 2; CF 2; CFG 2; BE 2; BEG 2; BCFG 2; BCDE 2; BEF 2; CDE 2; CDEG 1; BCDEG 1; ABCE 1; CDEF 1; EG 1; ACE 1; BCDEF 1; EF 1; C 1; BEFG 1; ACEG 1; EFG 1; ACEF 1; ABCEG 1; ABCEF 1; CDEFG 1; BCD 1; BCDG 1; BG 1; CDG 1; BCDF 1; E 1 |
| node 207 | BC 45; C 31; BCE 4; CE 3; BCG 2; BCF 2; BCD 2; ABC 2; CG 2; CF 2; CD 1; AC 1 |
| node 208 | BCE 9; CE 7; BCEG 6; CEG 5; BCEF 5; CEF 4; BCDE 4; CDE 3; BCEFG 3; CEFG 3; ABCE 2; BCDEG 2; BC 2; CDEG 2; BCDEF 2; BCG 2; ACE 2; CDEF 2; BCF 2; ABCEG 1; ACEG 1; CG 1; ABCEF 1; ACEF 1; CF 1; BCD 1; CDEFG 1; BCFG 1; ABCDE 1; CFG 1; BCDG 1; C 1; ACDE 1; BCDF 1; ABC 1; CDG 1; ACEFG 1; CD 1; CDF 1; ABCG 1; ACDEG 1; BE 1; BEG 1; ACDEF 1; ABCF 1 |
| node 209 | BCE 6; BCDE 5; CDE 5; CE 5; BCEG 4; CEG 4; BCDEG 4; CDEG 4; BCEF 4; CEF 4; BCDEF 3; CDEF 3; ABCE 3; ACE 3; BCEFG 3; ABCDE 3; CEFG 3; ACDE 2; ABCEG 2; ACEG 2; CDEFG 2; ABCEF 2; ACEF 2; ACDEG 2; ACDEF 2; ACEFG 1; BCD 1; BCDG 1; BCDF 1; BCG 1; CDG 1 |

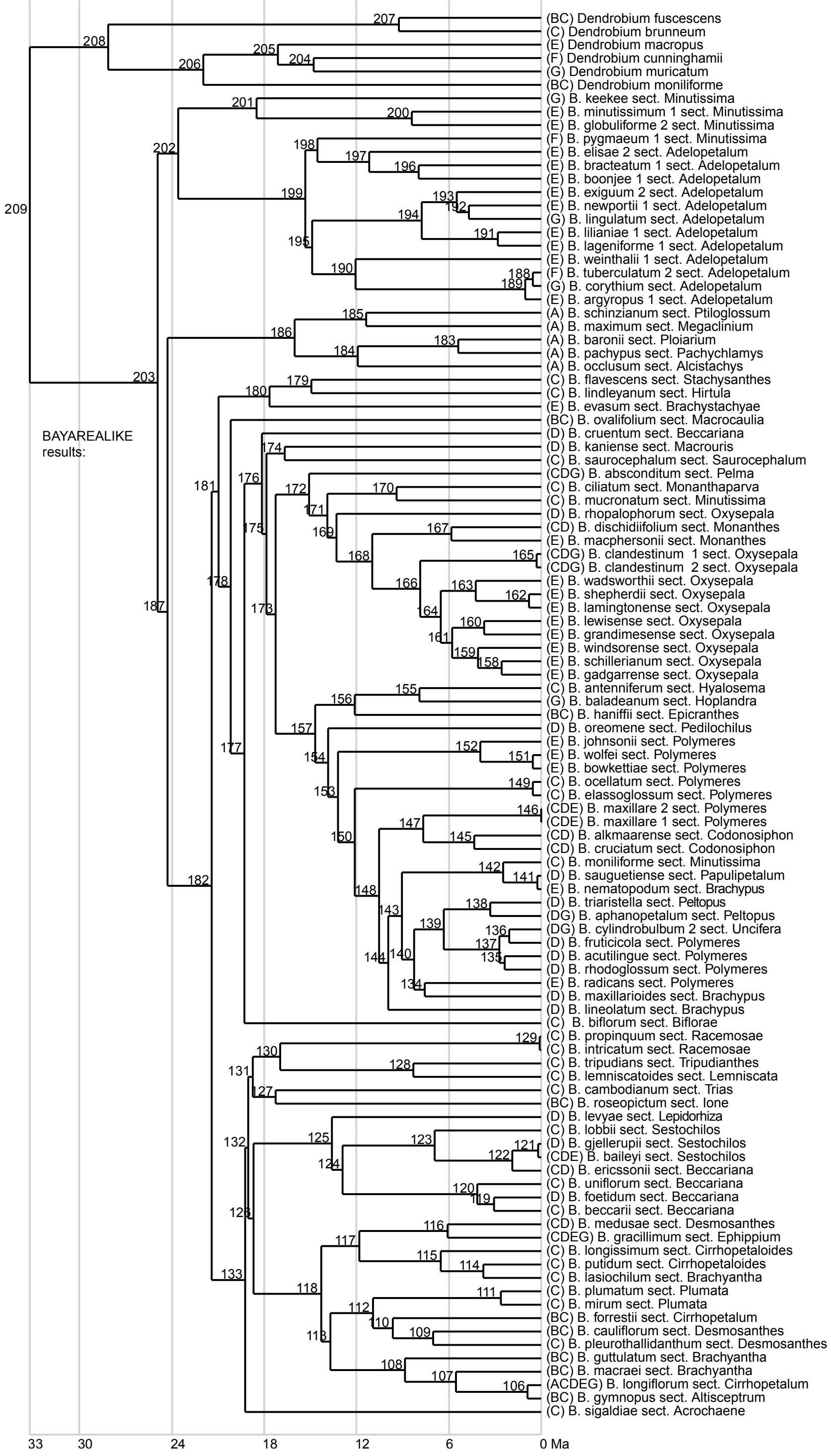


**Supplementary material S10: Nodes IDs for ancestral area reconstruction range probabilities.**
